# The role of syndapin-2 mediated transcytosis across the blood-brain barrier on amyloid-β accumulation in the brain

**DOI:** 10.1101/2020.07.12.199869

**Authors:** Diana M. Leite, Mohsen Seifi, Lorena Ruiz-Perez, Filomain Nguemo, Markus Plomann, Jerome D. Swinny, Giuseppe Battaglia

## Abstract

A deficient transport of amyloid-(Aβ) across the blood-brain barrier (BBB), and its diminished clearance from the brain, contributes to neurodegenerative and vascular pathologies, including Alzheimer’s (AD) and cerebral angiopathy, respectively. At the BBB, Aβ efflux transport is associated with the low-density receptor-related protein 1 (LRP1). However, the precise mechanisms governing Aβ transport across the BBB, in health and disease, remain to be fully understood. Recent evidence indicates that the LRP1 transcytosis occurs through a tubulation-mediated mechanism stabilised by syndapin-2. Here, we show that syndapin-2 is associated with Aβ clearance via LRP1 across the BBB. We further demonstrate that risk factors for AD, Aβ expression and ageing, are associated with a decline in the native expression of syndapin-2 within brain endothelium. Our data reveal that the syndapin-2-mediated pathway, and its balance with the endosomal sorting, are important for Aβ clearance proposing a measure to evaluate AD and ageing, as well as a target for counteracting Aβ build-up. Moreover, we provide evidence for the impact of the avidity of Aβ assemblies in their trafficking across the brain endothelium and in LRP1 expression levels, which may affect the overall clearance of Aβ across the BBB.

## Introduction

An estimated 50 million people worldwide are suffering of memory loss and other cognitive dysfunctions, collectively defined as dementia (1). The World Health Organisation has anticipated an increase of 10 million new cases every year (1). Up to 80% of these cases are associated with Alzheimer’s disease (AD), and more worryingly, AD and other dementia mortality have increased steadily in the last decade becoming one of the top leading causes of death worldwide. Among different factors, the most common AD feature is the abnormal accumulation of amyloid-β (Aβ) aggregates in the brain. Aβ is an end-product of the sequential processing of amyloid precursor protein (APP), and is implicated in a host of normal (2) and pathological functions within the brain parenchyma and blood vessels (3, 4). An essential process in the APP-Aβ pathway is the effective clearance of Aβ from the brain *via* the brain endothelium, also known as the blood-brain barrier (BBB), yet the exact mechanisms and molecular machinery remain to be fully elucidated. The importance of identifying how Aβ is transported from the brain to the blood is brought into sharp focus by the significant effect that impaired Aβ clearance has on the brain and vasculature health, over the course of lifetime. Indeed, an augmented expression of Aβ is a hallmark of ageing and age-related neurodegenerative disorders, including AD (5–8). Moreover, the accumulation of Aβ in the brain blood vessels contributes to cerebral amyloid angiopathy (9), also part of the AD pathology spectrum (10).

About 85% of all brain Aβ clearance occurs through the BBB (11), and it is expected that neurovascular dysfunctions contribute to defective Aβ clearance in AD (12, 13). The low-density lipoprotein receptor-related protein 1 (LRP1) is an essential receptor for the transport of brain Aβ across the BBB (11, 14–17). LRP1 is a multifunctional signalling and scavenger receptor consisting of a heavy chain that binds to various ligands, including apolipoprotein E (ApoE), *α*2-macroglobulin and APP (18). Importantly, a direct interaction of LRP1 and Aβ within the brain endothelium initiates clearance of Aβ from brain to blood through transcytosis (14). Other receptors, including glycoprotein 330 (19) and p-glycoprotein (20), as well as Aβ-binding proteins, such as ApoJ and ApoE (11, 21), regulate transport exchanges of their complexes with Aβ across the BBB.

Ageing is a prominent risk factor for the development of the sporadic form of AD, which accounts for ~ 95% of all cases (22, 23). Mounting evidence revealed that LRP1 expression declines in brain blood vessels and parenchyma during normal ageing in rodents and humans and is further reduced in AD individuals (11, 24–26). Moreover, validated genetic risk factors for AD, including ApoE E4 allele (27) and the gene encoding for phosphatidylinositol-binding clathrin assembly (PICALM) (28), are linked to diminished clearance of Aβ via LRP1. In the brain endothelium, Aβ binding to the ectodomain of LRP1 enhances the binding of PICALM that then initiates PICALM/clathrin dependent endocytosis of Aβ-LRP1 complexes through endocytic vesicles (such as, Rab5 and Rab11-positive endosomes) leading to Aβ transcytosis (28). Consequently, a reduction of the PICALM levels in AD impairs the mechanism of transcytosis of Aβ through LRP1/PICALM, and positively correlates with Aβ pathology and deterioration of cognition (28). In senile plaques, LRP1 ligands, such as ApoE, urokinase-type plasminogen activator and tissue plasminogen activator, co-deposit with Aβ indicating a loss of LRP1 in AD (29). Despite the amount of evidence demonstrating the fundamental role in the clearance of Aβ, how LRP1 controls transcytosis across the brain endothelium remains to be fully elucidated.

The relevance of Aβ-LRP1 trafficking in AD pathogenesis is accentuated by the identification of several endocytic-related genes that amplify the risk of late-onset AD, including PICALM, BIN1, and RIN3 (30–32), and abnormalities in the vesicular endosomal system (33–39). Hence, dysfunctions in the endocytic pathways appear to contribute to the AD pathology. Recently, we have demonstrated that at the brain endothelium, LRP1 associates with elements of classical vesicular endocytosis pathway (including clathrin, dynamin, and the early and late endosomes), as well as, with syndapin-2 (or PACSIN-2) to facilitate a fast tubulation-driven transcytosis (40). Syndapin-2 is a F-Bin/Amphiphysin/Rvs (F-BAR) protein, and due to the F-BAR domain, syndapin-2 senses and/or induces positive curvature (i.e., membrane bends in the direction of the leaflet decorated by the protein forming invaginations) stabilising tubular carriers (41). In addition, syndapin-2 contains a Src homology 3 (SH3) domain that binds to dynamin-2 and to actin-nucleating protein N-WASP that, ultimately, regulates actin filaments (42, 43). Our recent study unravelled that syndapin-2 is expressed in the brain blood vessels, and that syndapin-2 associates with LRP1 to drive a tubulation-mediated transcytosis (40). Importantly, one of the parameters that influence tubular transcytosis at the endothelium is how strong the cargo binds to its receptor, i.e., its affinity or avidity if multivalent (40, 44). By studying a model cargo targeting LRP1 decorated with different number of LRP1 ligands, we have revealed that a high number of ligands triggers internalisation through a conventional vesicular endocytic pathway leading to endo-lysosomal sorting and degradation, while a mid-avidity cargo undergoes transport across brain endothelium via syndapin-2-tubular carriers (40). We thus established that avidity enables high efficiency of transport across the BBB. Based on this work indicating the involvement of syndapin-2 in orchestrating LRP1 transcytosis, together with the recognised importance of LRP1 in the Aβ clearance from the brain, we hypothesised whether syndapin-2 is involved in the transport of Aβ and contributes to the build-up of Aβ in the brain. Furthermore, considering the influence of cargo avidity in the transport across the BBB, we evaluated whether different assemblies of Aβ, i.e., monomers, oligomers, and fibrils, trigger distinct mechanisms of internalisation and trafficking across the brain endothelium.

## Results

### Syndapin-2 directly interacts with LRP1 and Aβ-binding proteins

We first confirmed that syndapin-2 is endogenously expressed in brain endothelium using mouse brain endothelial cells (BECs). Immunoblotting of polarised BECs confirmed robust expression of syndapin-2 (**Fig. S1A**) while immunofluorescence showed that syndapin-2 immunoreactivity was localised to perinuclear vesicles and vesicular-tubular structures (**Fig. S1B**). These syndapin-2 positive vesicular-tubular structures exhibited an estimated diameter of 500 nm and lengths up to ~ 2 μm, which is in accordance to the tubules found for syndapin-2 on mouse brain capillaries images by stimulated emission depletion (STED) microscopy (40). Having confirmed the expression and location of syndapin-2 in BECs, we evaluated whether it is associated with receptors implicated in the transport of Aβ across the BBB. As shown in **Fig. 1A** and **Fig. S2A**, syndapin-2 is greatly associated with LRP1, while a much lower association was obtained for the receptor for advanced glycation end-products (RAGE) and p-glycoprotein (PGP), which are implicated in blood-to-brain and brain-to-blood transport of Aβ, respectively (20, 45). In a 3D rendering of a proximity ligation assay (PLA) of LRP1/syndapin-2, it was found that the proximity dots appear as long tubules spanning almost the entire cell thickness, whereas RAGE/syndapin-2 and PGP/syndapin-2 proximity dots resemble small spots (**Fig. S2B**). Having established the expression and location of syndapin-2 within BECs, and its association with LRP1 into tubular structures compared to other receptors involved in transport of Aβ across the BBB, we evaluated whether syndapin-2 interacts with LRP1 ligands, the Aβ-binding protein ApoE (**Fig. 1B-C**). Syndapin-2 is widely associated with LRP1, as evidenced by the abundance (~ 250 dots per cell) of PLA dots with each dot representing individual syndapin-2/LRP1 proteins located within 40 nm of another. Although, syndapin-2 was also located in a close proximity to ApoE (~ 250 dots per cell). Since ApoE is a LRP1 cognate ligand, and thus anticipated to interact with this receptor, we validated the PLA by quantifying the interaction of LRP1/ApoE (**Fig. 1C**). Indeed, we obtained a significant association between ApoE and LRP1 with substantial number of dots per cell (~ 80 dots). Interestingly, the abundance of dots for LRP1/ApoE was lower than that for syndapin-2/ApoE and syndapin-2/LRP1. Thus, these results suggest that syndapin-2 may also cooperate with other receptors for ApoE and, most importantly, other ligands for the LRP1. Syndapin-2 not only associates with the Aβ receptor (LRP1) but also ApoE, which might impact the clearance of Aβ across brain endothelium (**Fig. 1D**). Based on the *in vitro* findings, we then confirmed the expression of syndapin-2 in native mammalian brain. Immunoblotting of whole mouse brain homogenates demonstrated a strong syndapin-2 expression in the brain (**Fig. S1A**). Immunofluorescence revealed that syndapin-2 expression is enriched in the hippocampus, particularly in CA3 and the molecular layers of the dentate gyrus (**Fig. S1C**), as well as, in the molecular layers of the cerebellum (**Fig. S1D**), which is in agreement with previously published reports (46). The strong interaction of syndapin-2 and LRP1 in the brain was also confirmed in fixed mouse hippocampal tissue sections, by evaluating the colocalisation of their immunoreactivity profiles within brain blood vessels. Syndapin-2 and LRP1 immunoreactivity in brain blood vessels exhibited a strong colocalisation with a correlation coefficient value of 0.55 ± 0.28 (*n* = 40 blood vessels) (**Fig. 1E**). This correlation is clearly presented by the overlapping of syndapin-2 and LRP1 labelling on lectin-labelled blood vessels with syndapin-2 and LRP1 outlining the blood vessels within the hippocampus (**Fig. 1F)**. Our data demonstrates that in the brain blood vessels syndapin-2 associates with proteins involved in the machinery of Aβ clearance through the BBB, namely, LRP1 and ApoE.

**Figure 1.**
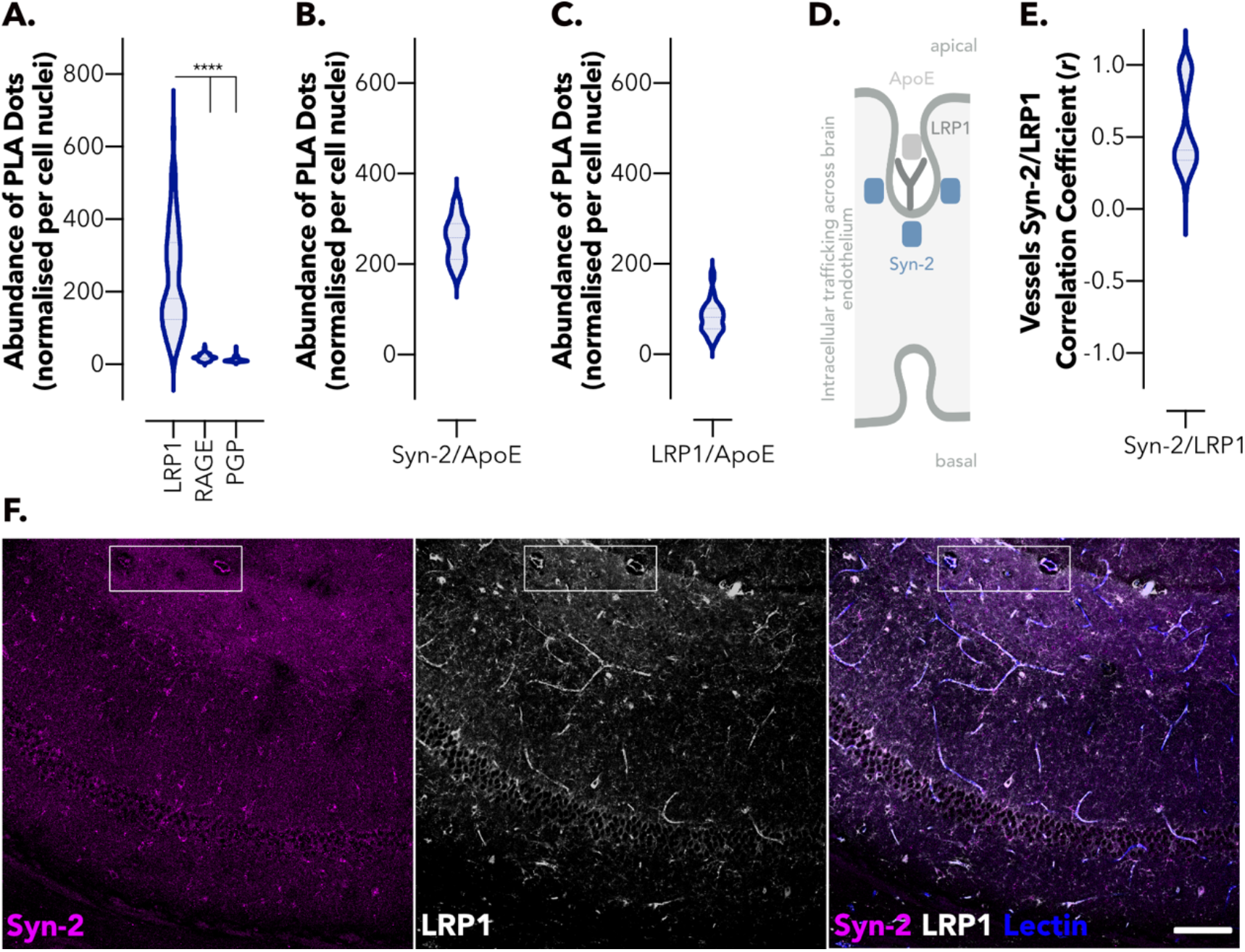
Syndapin-2 associates with LRP1 and Aβ-binding proteins within the brain endothelium. (**A**) Abundance of PLA dots resulting from the proximity of syndapin-2 with LRP1, RAGE, and PGP in BECs. Violin plot (*n* = 60 images). **** *P* < 0.0001, one-way ANOVA. Abundance of PLA dots resulting from the proximity of (**B**) syndapin-2 and ApoE and (**C**) LRP1 and ApoE in BECs. Violin plots (*n* = 20 to 40 images). (**D**) Schematic representation of the intracellular trafficking of LRP1/syndapin-2/ApoE complexes across BECs. (**E**) *Ex vivo* colocalisation of syndapin-2/LRP1 within the blood vessels of a mouse brain. Violin plot (*n* = 40 blood vessels). (**F**) Representative confocal images of the hippocampus of a mouse brain, indicating the immunoreactivity for syndapin-2 (in magenta) and LRP1 (in white) within lectin-labelled blood vessels in blue. The region highlighted with a box shows a region of blood vessels clearly showing colocalisation of syndapin-2 and LRP1 within the blood vessel walls.

### Vesicular and tubular LRP1 transcytosis in brain endothelium

Although transcytosis of LRP1 across the brain endothelium is well-established (40, 47–49), the precise mechanisms of the intracellular trafficking of LRP1 remains to be fully elucidated. In our recent study (40), we established that LRP1-mediated transcytosis occurs either through classical endocytic vesicles or tubular structures stabilised by syndapin-2. To further identify the cellular components associated with syndapin-2 tubular transcytosis, we performed a PLA between syndapin-2 and early-stage endocytosis effectors (heavy chain of clathrin and dynamin-2), early (EEA-1, Rab5), late (Rab7) and recycling endosomes (Rab11), lysosomes (LAMP1) and cytoskeletal elements (β-actin). Since it is well-established a role for PICALM in Aβ transcytosis via LRP1 (28), we also assessed syndapin-2/PICALM interaction in BECs. Our PLA analyses suggested negligible association of syndapin-2 with vesicular endocytic elements, as confirmed by the low number of PLA dots (< 20 dots per cell) resulting from association of syndapin-2 with early, late and recycling endosomes (**Fig. 2A**). Interestingly, a low association was also observed for syndapin-2 and PICALM, which may indicate that Aβ transcytosis mediated by PICALM is independent of the BAR protein. A stark contrast in the number of PLA dots was obtained for syndapin-2 and clathrin, dynamin-2 and β-actin (~ 150 dots per cell) compared to that of syndapin-2 and endosomal elements, as depicted in **Fig. 2A**. As previously reported (42, 43), syndapin-2 contains a SH3 domain that binds to dynamin-2 and N-WASP, which ultimately regulates actin filaments. Hence, this is in agreement with the abundant number of PLA dots obtained for syndapin-2 between dynamin-2 and β-actin. Syndapin-2 is generally associated with caveolae membrane sculpting (50) with little evidence showing a role for syndapin-2 in clathrin-mediated endocytosis. Our PLA results demonstrated a substantial interaction of syndapin-2 and clathrin, however it remains to be fully elucidated the involvement of clathrin in the mechanism of transcytosis mediated by syndapin-2. It is conceivable to suggest that syndapin-2-stabilised tubules are independent from the classical vesicular endocytosis. To further confirm if these two pathways operate wholly independently or they are capable of a compensatory crosstalk to counteract any dysregulation that may arise in one another, we evaluated whether the downregulation of syndapin-2 affects the intracellular trafficking of LRP1. To do so, we established BECs expressing low levels of syndapin-2, by knocking down the expression of syndapin-2 using a short hairpin RNA (shRNA) (40), and assessed the association of LRP1 with cellular components involved in transcytosis. It is worth to highlight that syndapin-2 knockdown in BECs resulted in no significant effect on the BBB properties with monolayers exhibiting dextran permeability values of 25.6 and 5.3 nm s^−1^ for 4 and 70 kDa dextrans, respectively (40). We also confirmed that LRP1 expression remains unaltered with the knock-down of syndapin-2 (**Fig. 2B**). Knockdown of syndapin-2 in BECs triggered in a significant reduction (∼ 2-fold change) in the abundance of PLA dots resulting from LRP1/syndapin-2 (**Fig. 2C**). In contrast, a downregulation of syndapin-2 led to an increase in the association of LRP1 with heavy-chain of clathrin (**Fig. 2D**). This increase in LRP1/syndapin-2 dots may suggest that the mechanism of trafficking by syndapin-2 is independent from clathrin recruitment to the membrane for the formation of clathrin-coated pits, as more clathrin is in proximity to LRP1 with a reduction in the expression levels of syndapin-2. Notably, downregulation of syndapin-2 caused an increase in the association of LRP1 with early endosomal proteins (EEA-1, Rab5) (**Fig. 2E-F**), as well as late endosomes (Rab7) (~ 2-fold change in PLA dots) (**Fig. 2G**). Other stages of endocytosis including recycling endosomes (Rab11) and lysosomal degradation (LAMP-1), appeared to be not affected by the levels of syndapin-2 in BECs (**Fig. 2H-I**). Thus, a depletion of syndapin-2 levels generates a substantial increase in the vesicular trafficking of LRP1 with the receptor interacting with early endocytic effectors and early and late endosomal markers (EEA-1, Rab5, Rab7). To investigate whether the depletion of syndapin-2 triggers alterations in the vesicular endosomal pathway, we evaluated the brain blood vessels of heterozygous syndapin-2 knockout mice (KO) compared to wild-type (WT). Microvessels from WT and KO mouse brains were separated by a gradient centrifugation method performed on whole brains (28, 51) in which two fractions are obtained containing the microvessels and parenchymal cells. Initially, we analysed the LRP1 levels at the microvessels of WT and heterozygous KO mouse brains (**Fig. 2J, L**). No significant alterations were observed in the expression levels of LRP1, which also corroborates the *in vitro* results obtained in syndapin-2 knockdown BECs. We then measured the levels of the early endocytic protein Rab5, which we found to greatly associate with LRP1 once syndapin-2 expression is reduced in BECs. Notably, we found that the heterozygous KO microvessels expressed a higher level of Rab5 than the ones of the WT (**Fig. 2K, L**). This suggests that a depletion of syndapin-2 triggers a compensatory mechanism in intracellular trafficking by increasing early endocytic proteins. Under basal conditions, syndapin-2-mediated LRP1 trafficking is independent from vesicular endocytic trafficking. However, alterations in syndapin-2 levels pivots LRP1 trafficking towards a vesicular pathway resulting in an increase in Rab5 levels as a compensatory mechanism thereby emphasising the importance of syndapin-2 in determining transport in BECs.

**Figure 2.**
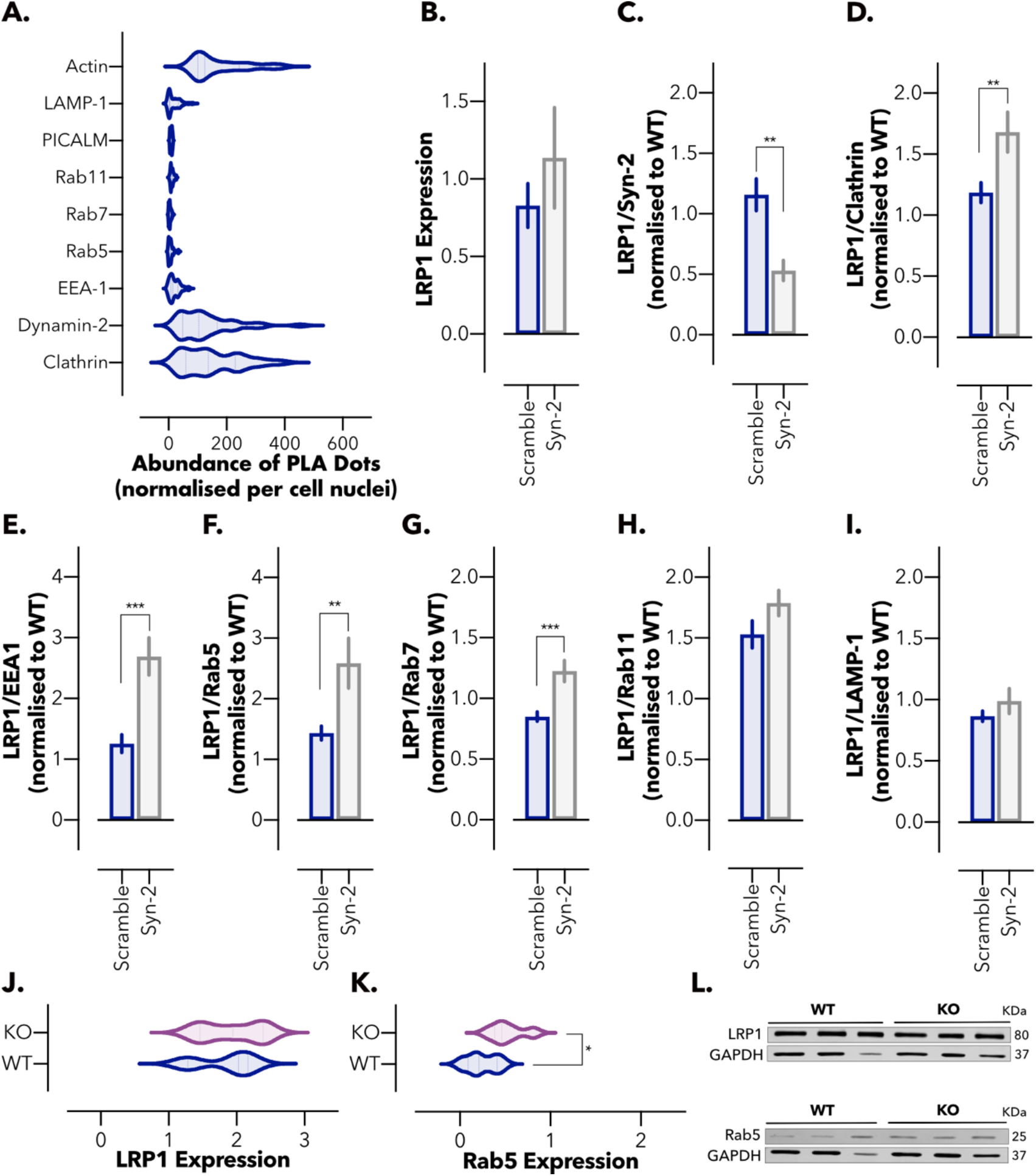
Balance between the vesicular and tubular trafficking of LRP1 in the brain endothelium. (**A**) Abundance of PLA dots resulting from the proximity between syndapin-2 and intracellular proteins associated with endocytosis and trafficking of LRP1 and Aβ in BECs. Violin plot (*n* = 30-40 images). (**B**) Expression levels of LRP1 in the shRNA control (scramble) and syndapin-2 knockdown (Syn-2) BECs. Values normalised to wild-type BECs. GAPDH was used as loading control. Mean ± SD (*n* = 3). Non-significant, Student’s t-test. Abundance of PLA dots resulting from the proximity between LRP1 and (**C**) syndapin-2, (**D**) clathrin, (**E**) EEA-1, (**F**) Rab5, (**G**) Rab7, (**H**) Rab11, and (**I**) LAMP-1 in scramble and syndapin-2 knockdown BECs. All data were normalised to wild-type (WT) BECs. Mean ± SD (*n* = 30 to 40 images). ** *P* < 0.01, *** *P* < 0.001, Student’s t-test. (**J**) LRP1 and (**K**) Rab5 expression levels in the microvessels of wild-type (WT) and heterozygous syndapin-2 knockout (KO) mouse brains quantified by densiometry analyses of immunoblotting. Data normalised to GAPDH (loading control). Violin plot (*n* = 3-5). * *P* < 0.05, Student’s t-test. (**L**) Representative immunoblotting for syndapin-2, Rab5 and GAPDH (as loading control) in the microvessel fractions of WT and KO mouse brains.

### Syndapin-2 interacts with Aβ and transports it across the brain endothelium

Given the role of syndapin-2 in mediating the LRP1 trafficking and the recognised importance of LRP1 in Aβ clearance across the brain endothelium (28), we next evaluated whether syndapin-2 is involved in the transport of Aβ across BECs. Polarised BECs were incubated with FAM-Aβ (1-40) (500 nM) for 5 minutes, and the colocalisation levels of FAM-Aβ (1-40) and syndapin-2 were quantified (**Fig. 3A-C**). In **Fig. 3A**, Aβ (1-40) was located within the cell cytoplasm in close proximity to the immunoreactivity for syndapin-2. Horizontal orthogonal views (*x, z*) obtained from *z*-stacks of the *in vitro* monolayer of BECs exhibited that colocalised syndapin-2/Aβ (1-40) profiles within the cells were presented as elongated tubular-like structures (**Fig. 3B**). Importantly, these syndapin-2/Aβ complexes appeared not only at the apical and basal membranes but also within the cell, most likely *en route* from the basal to apical side of the BECs. This implies that syndapin-2 remains complexed with Aβ endocytic tubular structures in early and late stages of endocytosis and, possibly, acting together with cytoskeletal proteins, such as actin (43) (**Fig. 2A**), to modulate deformations of the cell membrane into tubules. To further investigate the interaction of syndapin-2 and FAM-Aβ, we analysed their degree of colocalisation in polarised BECs (**Fig.3C**). The correlation coefficient of ∼ 0.4 corroborated the strong interaction of syndapin-2 with Aβ. Since other receptors, including RAGE and PGP, might be involved in the transport of Aβ in BECs, we assessed whether the presence of FAM-Aβ (1-40) triggered an association of syndapin-2 with these receptors. As shown in **Fig. S3A**, the addition of FAM-Aβ (1-40) for 15 minutes caused a significant increase in the abundance of PLA dots of LRP1/syndapin-2, while no significant difference was observed for the interaction of syndapin-2 with RAGE or PGP. Therefore, our findings suggest that the syndapin-2/Aβ (1-40) complexes trafficking relies on the recruitment of syndapin-2 to the cell membrane upon binding of Aβ (1-40) to LRP1. To evaluate whether the association of syndapin-2 and Aβ holds true in the mammalian brain, we studied the colocalisation of syndapin-2 and Aβ immunoreactivity in coronal brain sections of a wild-type mouse hippocampus (**Fig. 3D**). 3D reconstructions from regions of interest (ROI) clearly demonstrated a close association between syndapin-2 and Aβ within the lectin-labelled blood vessels. To further confirm the involvement of syndapin-2 in Aβ transport, we investigated whether depletion of syndapin-2 in BECs influences the permeability of Aβ across our *in vitro* BBB model in the basal-to-apical direction (brain-to-blood). Using syndapin-2 knockdown BECs, we found that a depletion in the expression levels of syndapin-2 caused a significant reduction in the basal-to-apical permeability of FAM-Aβ (1-40) (**Fig. 3E**). It is worth to mention that, in the absence of syndapin-2, BECs triggered the vesicular endosomal pathway for the intracellular trafficking of LRP1 (**Fig. 2D-J**). Thus, it is conceivable to hypothesise that in syndapin-2 knockdown BECs, LRP1/Aβ (1-40) complexes are transported through a vesicular endosomal pathway to compensate the depletion of tubular-mediated transcytosis stabilised by syndapin-2. Nevertheless, a reduction in syndapin-2 expression levels still impaired the transport of Aβ (by ~ 20% compared to BECs expressing endogenous levels). Interestingly, we did not observe a significant alteration in the apical-to-basal transport of FAM-Aβ (1-40) in BECs lacking syndapin-2 (**Fig. S3B**). RAGE transports Aβ across the BECs from the blood to the brain (45). As the addition of FAM-Aβ (1-40) resulted in no significant effect in the association of syndapin-2/RAGE (**Fig. S3A**), it may be that the Aβ transport through RAGE is not affected by a depletion in the expression levels of syndapin-2 and, consequently, no differences observed in the apical-to-basal transport of Aβ. To further establish the role of syndapin-2 in the accumulation of Aβ in the brain, we quantified the amount of Aβ (1-40) in the brain and plasma of syndapin-2 KO mice and compared to age-matched WT using ELISA. As depicted in **Fig. 3F**, the levels of Aβ (1-40) in the brains of heterozygous KO mice were significantly higher than in the WT brains, while statistically significant difference was obtained for Aβ (1-40) levels in the plasma (**Fig. 3G**). These results corroborate that depletion of syndapin-2 contributes to a deficient transport of Aβ across the BBB, which consequently leads to accumulation of Aβ in the brain. Collectively, these data provide the first demonstration for the implication of syndapin-2 in Aβ trafficking across BECs and accumulation in the brain.

**Figure 3.**
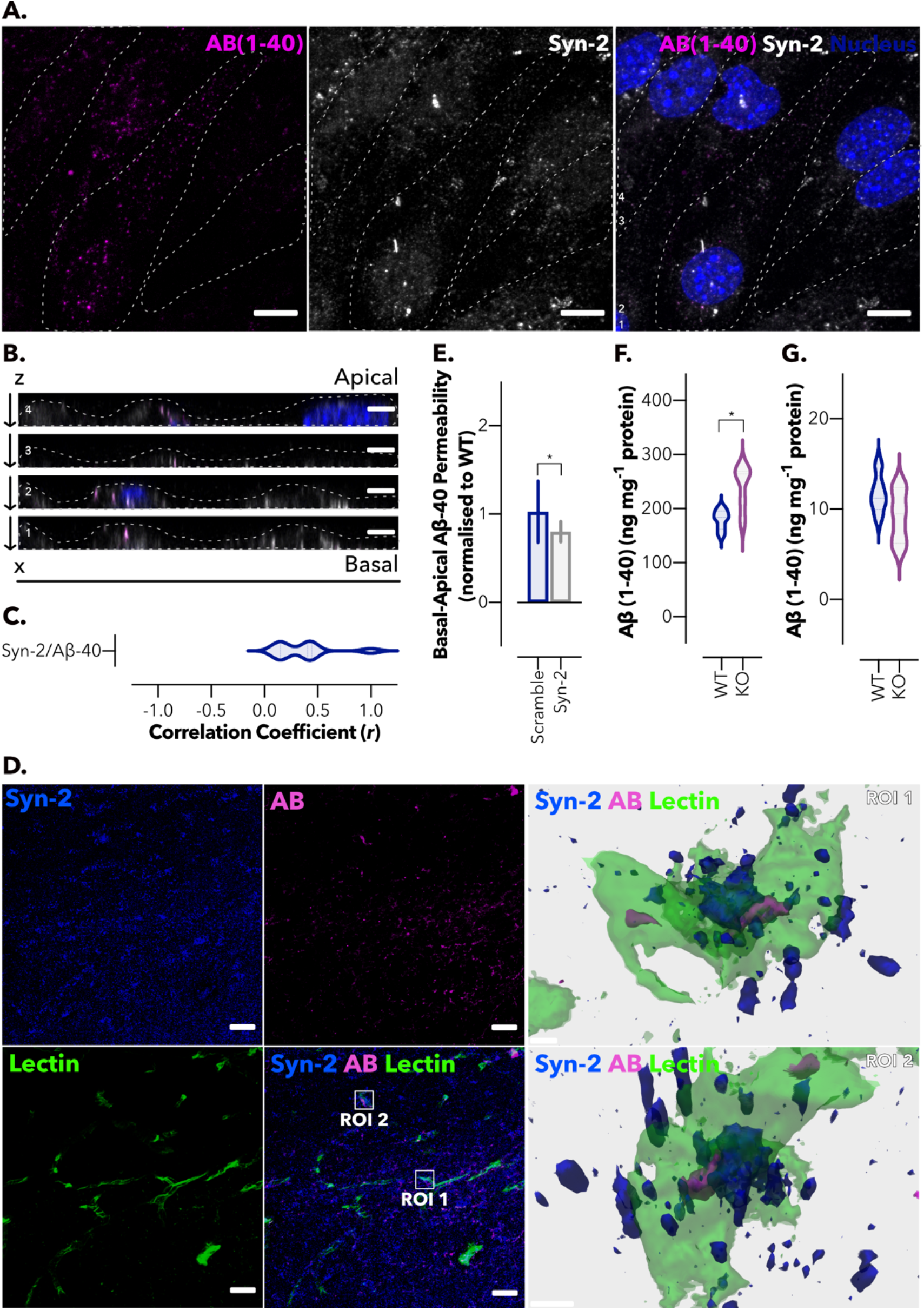
Syndapin-2 transports Aβ across the brain endothelium. (**A**) *In vitro* colocalisation of Aβ (1-40) (in magenta) and syndapin-2 (in white) in polarised BECs treated with FAM-Aβ (1-40) for 5 minutes. Nuclei are shown in blue. Dotted line represents the cell membrane limits. Scale bar: 10 μm. (**B**) Z-projections of BECs showing the colocalisation of syndapin-2 with Aβ (1-40) during basal-to-apical transport. Dotted lines represent the cell membrane limits. Scale bar: 5 μm. (**C**) Quantification of colocalisation of syndapin-2 and Aβ (1-40) within the polarised BECs. Violin plot (*n* = 10 images). (**D**) Representative confocal image of the hippocampal region of a mouse brain showing syndapin-2 (in blue), Aβ (in magenta) and lectin-labelled blood vessels (in green). Scale bar: 20 μm. 3D reconstructions of a z-stack of two regions of interest (ROI) showing close interaction of syndapin-2 and Aβ in the brain blood vessels in the left panel. Scale bar: 5 μm. (**E**) *In vitro* permeability of FAM-Aβ (1-40) across polarised shRNA control (scramble) and syndapin-2 knockdown (Syn-2) in a basal-to-apical (brain-to-blood) direction. Mean ± SD (*n* = 15). * *P* < 0.05, Student’s t-test. Permeability values were normalised to wild-type (WT) BECs. Quantification of Aβ (1-40) in the (**F**) brain and (**G**) plasma of wild-type (WT) and syndapin-2 knockdown (KO) animals. Mean ± SD (*n* = 5-6). * *P* < 0.05, Student’s t-test.

### Syndapin-2 expression is reduced in the APP-PS1 mouse model of AD

Given that the downregulation of syndapin-2 expression levels in BECs resulted in a reduced brain-to-blood Aβ transport (**Fig. 3E**) and accumulation in the brain (**Fig. 3F**), we probed the interdependence of this relationship by measuring the impact of increased native Aβ expression on syndapin-2 levels in a mammalian brain using the APP-PS1 model. APP-PS1 mice accumulate Aβ at a young age and develop extracellular plaques consisting of fibrillary Aβ deposits, in the cerebral cortex and hippocampus, commencing at around 6 months of age (52), and confirmed in our mouse line, using 12-month old subjects (**Fig. 4A**). Immunoreactivity for Aβ was distributed in the brain parenchyma and blood vessels, with signal for syndapin-2 localised to the wall of the blood vessels and the surrounding intravascular Aβ deposits (**Fig. 4A, ROI1**). Additionally, as observed in the WT littermates (**Fig. 3D**), syndapin-2 immunoreactivity was located on the basal surface of the blood vessels in close association with extravascular Aβ in the APP-PS1 brains (**Fig. 4A, ROI2**). This indicated that syndapin-2 is expressed in both WT and APP-PS1 brain, in association with Aβ, possibly facilitating its transport across the brain endothelium. To investigate the potential correlation between Aβ accumulation and alterations in vesicular and tubular endocytosis in the brain blood vessels, microvessels from WT and APP-PS1 mouse brains were separated by a gradient centrifugation method performed on whole brains (28, 51). Initially, in the microvessel fractions, the expression of LRP1, tubular-associated and vesicular endocytic proteins was assessed (**Fig. 4B-H**) (**Fig. S4A**). Immunoblotting densiometry analysis exhibited a slight decrease in the LRP1 expression levels in microvessels of APP-PS1 brains (**Fig. 4B**). We further studied alterations in the vesicular endosomal pathway. Notably, a significant increase (a ~ 3-fold change) was found for the levels of clathrin (**Fig. 4C**) and early endosomes, Rab5 (**Fig. 4D**), in APP-PS1 microvessels compared to that of WT littermates. Other endosomal proteins, namely, late (Rab7; **Fig. 4E**), recycling endosomes (Rab11; **Fig. 4F**), and lysosomes (LAMP-1; **Fig. 4G**) remained unaltered in the APP-PS1 brains compared to the WT littermates. Along with these alterations in the vesicular endosomal trafficking, we also found that syndapin-2 levels are reduced in the microvessels of APP-PS1 mouse brains compared to WT (**Fig. 4H**). This significant increase in vesicular endosomal proteins suggests the balance between vesicular and tubular transport with the intracellular trafficking shifting dynamically depending on the syndapin-2 expression levels within the brain endothelium. Since syndapin-2 levels in isolated microvessels were comparable to those in the parenchymal fraction, which contained neurons and glia, we also measured the level of endocytic proteins in the parenchyma of WT and APP-PS1 brains (**Fig. S4B**). In the parenchymal fraction, LRP1 levels were found to be unaltered, while syndapin-2 levels were reduced in APP-PS1 parenchyma compared to WT brains. Along with the alteration in the syndapin-2 levels, an overexpression of Rab5 was found in the parenchymal fraction of APP-PS1, which confirms the previously reported overactivation of Rab5-positive endosomes in neurons of AD individuals (34, 35). Other endocytic proteins, including clathrin, Rab7 and Rab11, were found to remain unaffected in the APP-PS1 mouse brains compared to WT. Interestingly, an increase in the lysosomal marker was observed in the parenchymal fraction of APP-PS1 brains. Similar to what occurs at the microvessels, syndapin-2 might be involved in the overactivation of early endosomes in parenchymal cells. These findings also suggest a role for syndapin-2 in the trafficking of Aβ in neuronal and/or glial cells, in which dysfunctional syndapin-2 levels are accompanied by an increased number of early endosomes. Overall, our data provide the first evidence for the interaction of syndapin-2 with Aβ in the mammalian brain, and its dysfunction in an AD mouse model. Importantly, the syndapin-2 and Rab5 levels in younger animals (4-months old) were assessed within the microvessels and parenchyma (**Fig. S5**). However, no significant changes were observed between WT and APP-PS1 blood vessels. Hence, this indicates that alterations in tubular and vesicular endosomal transport at the BECs are age dependent.

**Figure 4.**
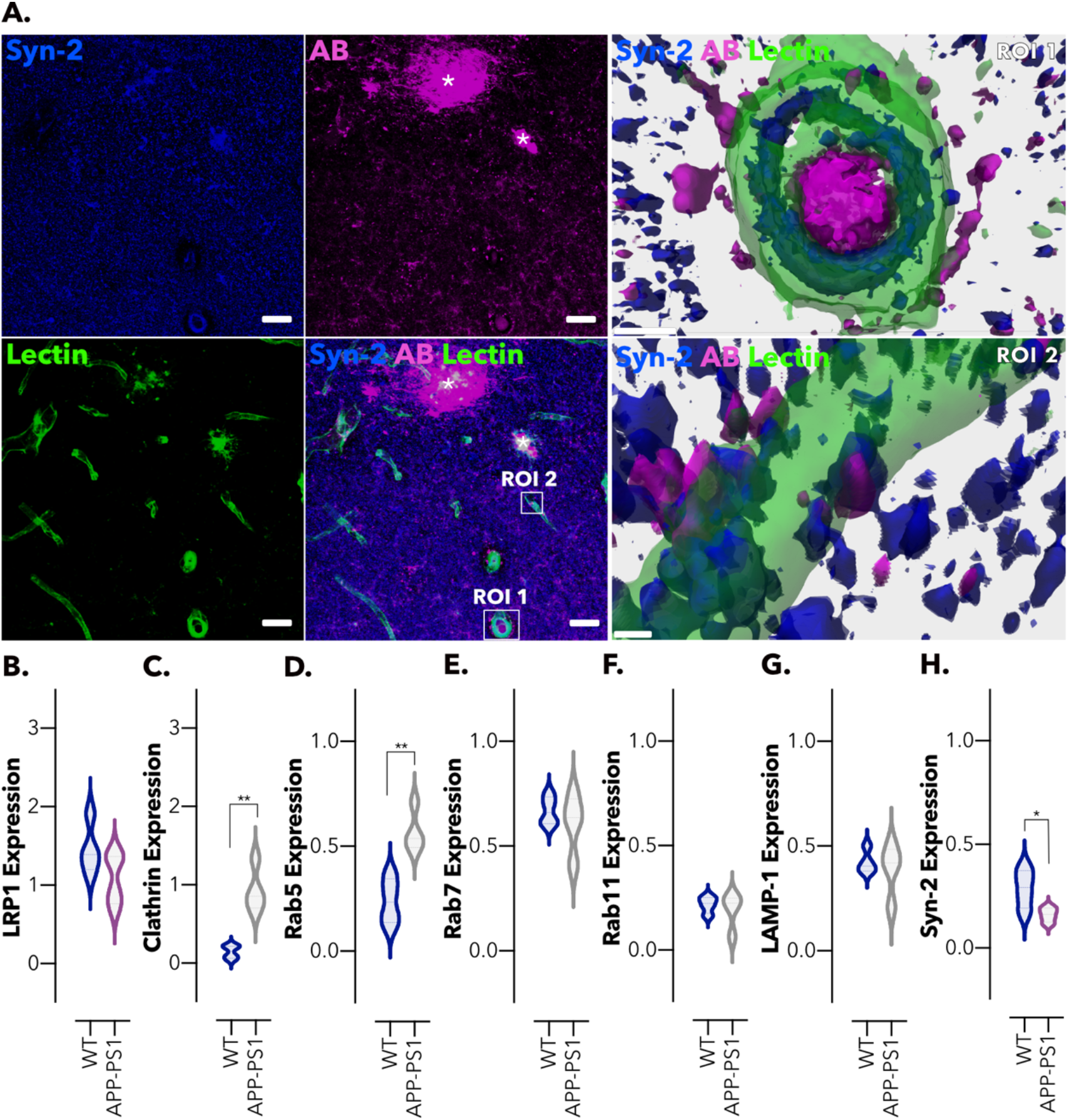
Intracellular trafficking is altered in brain microvessels in an amyloidosis mouse model. (**A**) Representative confocal images of the hippocampal region of an AD mouse brain (APP-PS1) showing syndapin-2 (in blue), Aβ (in magenta) with lectin-labelled blood vessels (in green). Stars denote Aβ plaques colocalising with syndapin-2. Scale bar: 20 μm. 3D reconstructions of z-stack acquired on two regions of interest (ROI1 and ROI2) within the hippocampus of an APP-PS1 mouse brain exhibiting the close association of syndapin-2 and Aβ within the blood vessels. Scale bar: 5 μm. Expression levels of (**B**) LRP1, (**C**) clathrin, (**D**) Rab5, (**E**) Rab7, (**F**) Rab11, (**G**) LAMP-1 and (**H**) syndapin-2 in the microvessels from WT and APP-PS1 mouse brains. Data normalised to GAPDH (loading control). Mean ± SD (*n* = 4 animals). * *P* < 0.05, ** *P* <0.01, Student’s t-test.

### Syndapin-2 expression levels are decreased with age

Ageing is the most established risk factor for developing AD, with sporadic AD accounting for ~ 95% of all cases (22, 23, 53). Within the blood vessels, the expression of LRP1, an efflux transporter for Aβ, declines in normal ageing resulting in an accumulation of Aβ (25). Moreover, genes encoding proteins which interact with LRP1 and that are involved in Aβ transcytosis across the brain endothelium, including PICALM and ApoE, have been identified as prominent risk factors for late-onset sporadic AD (54). Given the interaction of syndapin-2 with LRP1 and ApoE, we addressed whether the syndapin-2 levels are altered by healthy ageing. We thus evaluated syndapin-2 and Rab5 expression in microvessels and parenchyma of 4- and 12-months old WT animals, respectively (**Fig. 5A1-3**) (**Fig. S6A**). Immunoblotting densiometry analysis revealed that syndapin-2 expression was significantly reduced in the microvessels of 12-months old brains compared to 4-months old WT mice (**Fig. 5A2**) while GTPase Rab5 expression showed no significant alterations (**Fig. 5A3**). Consequently, this reduction in syndapin-2 levels resulted in a decrease of a syndapin-2/Rab5 ratio in the microvessels of healthy aged animals (**Fig. 5B**). In the parenchyma of WT mouse brains, no significant alterations were found in syndapin-2 and Rab5 expression levels (**Fig. S6A**). To determine whether AD risk factors of age and genetic mutations resulting in increased Aβ are additive to their influence on syndapin-2 and Rab5 expression, we also performed quantitative immunoblotting on brains obtained from 4- and 12-months old APP-PS1 mice (**Fig. 5C1-3**) (**Fig. S6B**). In the APP-PS1 brains, the expression levels of syndapin-2 were significantly reduced (by 6-fold) within the microvessels (**Fig. 5C2**), while Rab5 levels were significantly increased in 12-months old compared to 4-months old APP-PS1 brains (**Fig. 5C3**). Consistent with these results, the syndapin-2/Rab5 expression ratio in AD was appreciably decreased in 12-months old brains (**Fig. 5D**). Notably, the decline in this syndapin-2/Rab5 ratio within the microvessels in 12-months old APP-PS1 brains was much greater than that of WT. A similar trend was observed for the parenchyma fractions of APP-PS1 brains (**Fig. S6A-B**). However, a decline in syndapin-2 expression is more prominent in the microvessels. Hence, although the loss of syndapin-2 occurs naturally through the process of ageing, a genetic predisposition to AD appears to accelerate this loss. Our results provide the first evidence for the role of syndapin-2 in Aβ clearance from the brain through LRP1-mediated transcytosis across the brain endothelium, and that the process of healthy ageing also leads to a decline in the expression of syndapin-2, which is accelerated in AD.

**Figure 5.**
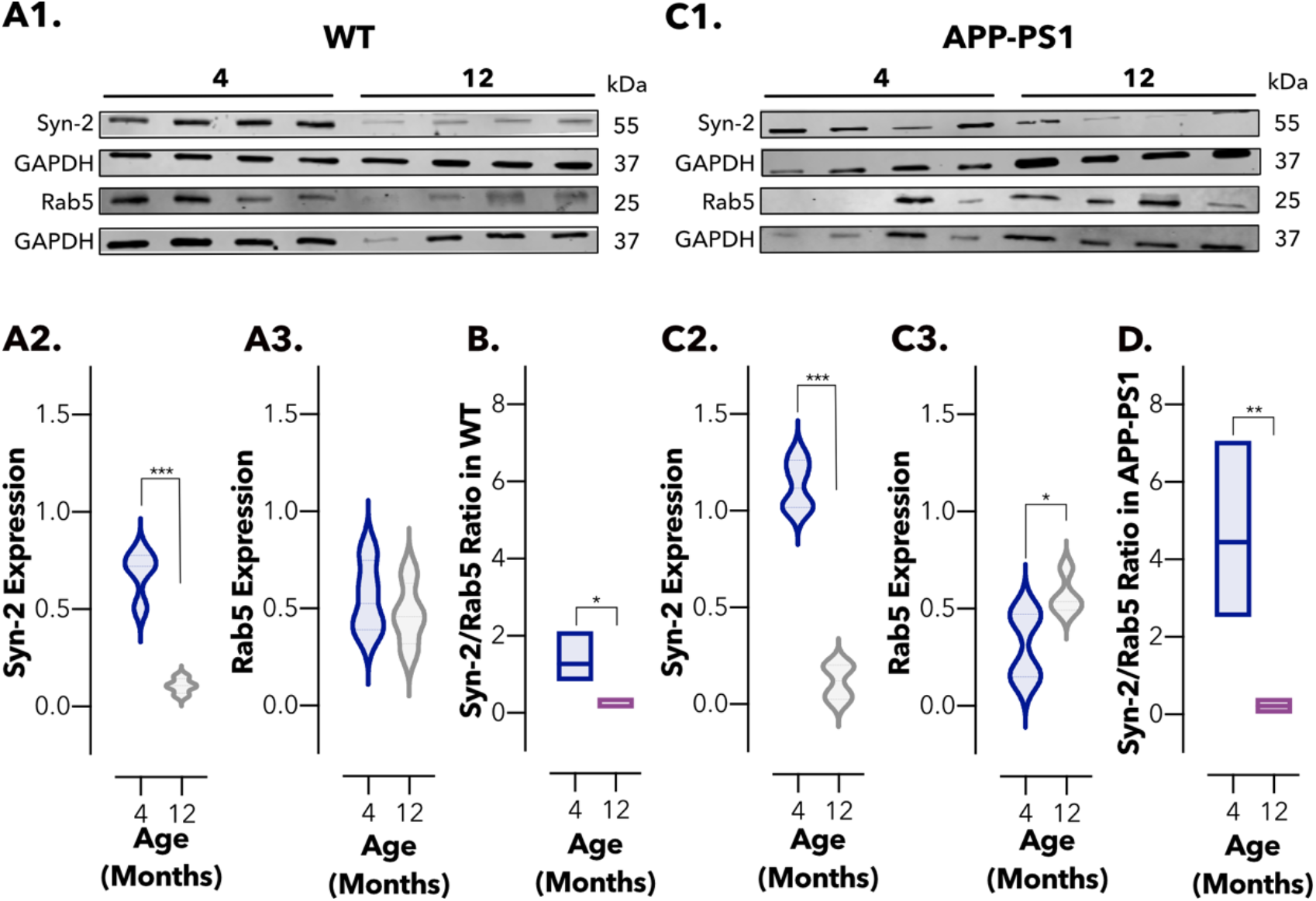
Imbalance in the levels of syndapin-2 in brain microvessels is associated with ageing. (**A1**) Immunoblotting for syndapin-2, Rab5 and GAPDH (as loading control) in the microvessel fractions of 4- and 12-months old WT mouse brains. Expression of (**A2**) syndapin-2 and (**A3**) Rab5 in microvessels of WT mouse brains. Data normalised to GAPDH. Violin plot of *n* = 4 animals. *** *P* <0.001, Student’s t-test. (**B**) Ratio of syndapin-2/Rab5 in WT brain microvessels. * *P* <0.05, Student’s t-test. (**B1**) Immunoblotting for syndapin-2, Rab5 and GAPDH (loading control) in the microvessels of 4- and 12-months old APP-PS1 mouse brains. Expression of (**C2**) syndapin-2 and (**C3**) Rab5 in the APP-PS1 brain microvessels. Data normalised to GAPDH. Violin plot of *n* = 4 animals. * *P* <0.05, *** *P* <0.001, Student’s t-test. (**D**) Ratio of syndapin-2/Rab5 in APP-PS1 microvessels. ** *P* <0.01, Student’s t-test.

### Avidity of Aβ assemblies dictates intracellular trafficking across brain endothelium

Aβ aggregates into different assemblies ranging from soluble oligomers to insoluble fibrils, and several studies have shown that oligomeric and fibrillar species of Aβ contribute differently to initiation, seeding and propagation of the AD pathology (55–57). As Aβ monomers assemble into oligomers and fibrils, the number of ligands that bind to LRP1 increases and with this the total avidity. Similar to the model cargo targeting LRP1 described in our previous study (40), we hypothesised that the Aβ distinct avidities toward LRP1 affect their intracellular trafficking and clearance across the BBB. We prepared monomeric, oligomeric and fibrillar Aβ assemblies (58, 59), and characterised these assemblies by TEM (**Fig. 6A**). Imaging of monomeric Aβ showed the presence of small assemblies and agglomerates (**Fig. 6A1, RO1 and RO2**) with no defined structures. In the oligomeric Aβ, populations of 5-15 nm globular structures were obtained (**Fig. 6A2, RO3**), in agreement with the morphologies reported in previous studies (58), while the fibrillar Aβ imaging exhibited the presence of fibrils (thickness of ~10 nm) coexisting with a few oligomers of ~ 15 nm (**Fig. 6A3**). Notably, populations of elongated pearl-necklace assemblies were also found in the fibrillar Aβ sample, in which it appears that fibrils are emerging from the oligomers (**Fig. 6A3, RO4**). Fibril structure was further confirmed by a thioflavin T (ThT) assay (**Fig. S7A**) with a characteristic increase in the fluorescence intensity of ThT after incubation with Aβ fibrils. We initially assessed the toxicity in BECs and, at the concentrations tested, none of the Aβ assemblies caused significant toxicity (**Fig. S7B**). We thus treated BECs with Aβ monomers, oligomers, and fibrils and assessed the interaction between Aβ and LRP1 with both tubular (syndapin-2) and vesicular (Rab5) endocytic markers (**Fig. 6B1, B2**) using a PLA. All assemblies caused an increase in LRP1/syndapin-2 association in comparison to untreated cells (> 2-fold increase), indicating that syndapin-2 is recruited when BECs are exposed to Aβ (**Fig. 6B1**). However, the oligomeric species caused the highest increase in LRP1/syndapin-2 interaction alongside with a decrease in LRP1/Rab5. Conversely, both monomers and fibrils triggered BECs to favour LRP1/Rab5 association (**Fig. 6B1, B2**). By tracking Aβ, we observed that Aβ oligomers are more associated with syndapin-2 (**Fig. 6C1**), while the monomers and fibrils are more in proximity to Rab5 (**Fig. 6C2**). As established in our previous work (40), association with syndapin-2 corresponds to fast transcytosis and tubulation, while association with Rab5 indicates endosomes sorting followed by degradation. We thus assessed, LRP1 expression levels following incubation with Aβ assemblies (**Fig. 6D**). Western blot analyses displayed that LRP1 expression is unaltered after incubation with monomers and oligomers (100 nM) for 2 hours. In contrast, incubation with Aβ fibrils (100 nM) resulted in a reduction in the expression levels of LRP1 (a 2-fold reduction). Collectively, our data demonstrates that Aβ oligomers (i.e., mid-avidity to LRP1) are sorted into syndapin-2 rich compartment and thus shuttled across, while fibrils (i.e., high avidity to LRP1) skew the trafficking toward endosomal sorting and degradation.

**Figure 6.**
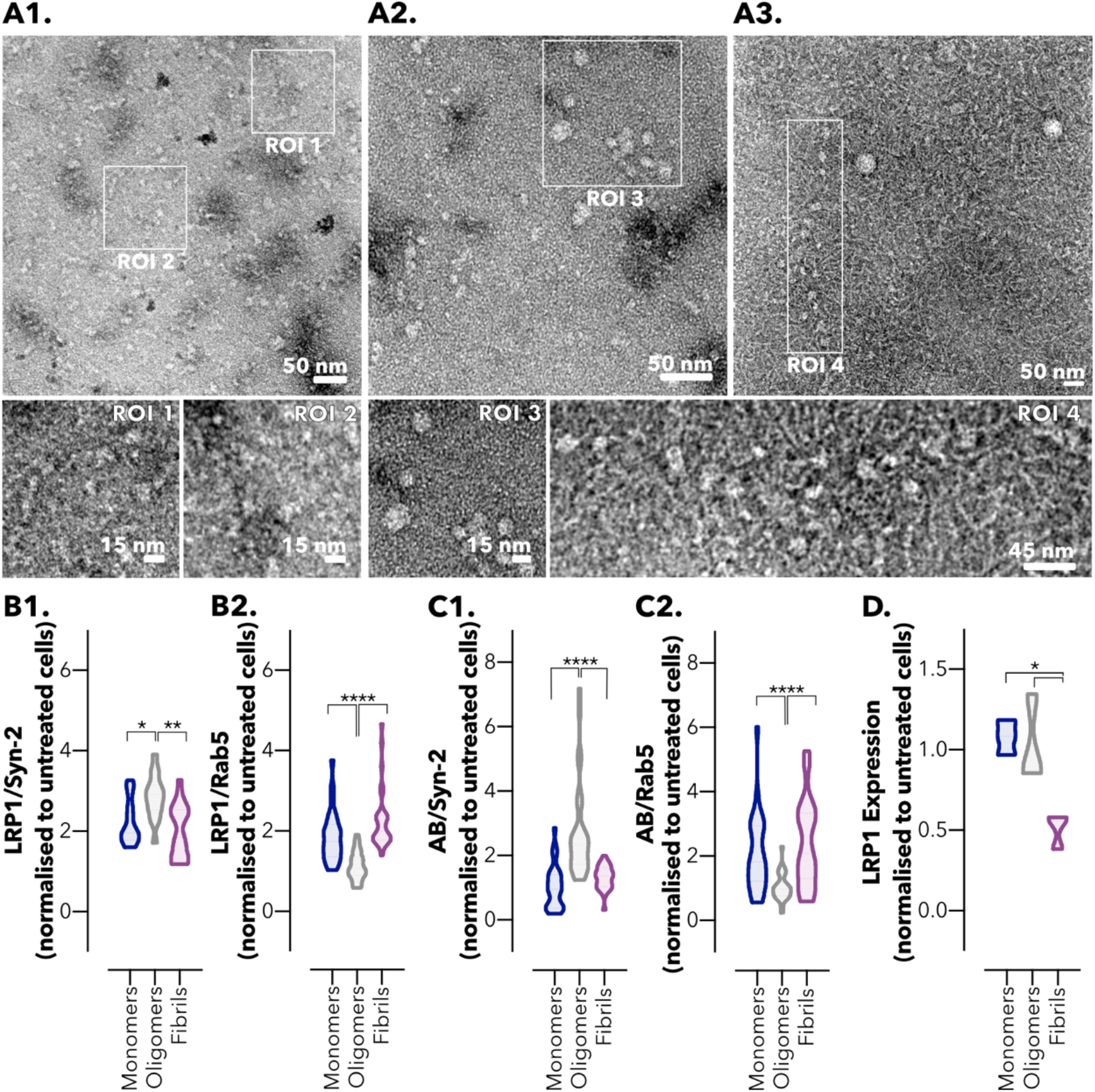
Avidity of Aβ assemblies dictates intracellular trafficking across the brain endothelium. Transmission electron micrographs of Aβ (**A1**) monomers, (**A2**) oligomers, and (**A3**) fibrils. Regions of interest (ROI) 1 and 2 denote details of the monomeric Aβ. ROI3 highlights oligomers and ROI4 shows a region of Aβ oligomers and fibrils. Abundance of PLA dots resulting from the proximity of LRP1 with (**B1**) syndapin-2 and (**B2**) Rab5 in BECs treated with Aβ assemblies (monomers, oligomers, and fibrils) at 100 nM for 15 minutes. Violin plot of *n* = 30 images. * *P* <0.05, ** *P* <0.01, **** *P* <0.0001, one-way ANOVA comparing oligomers *versus* monomers and fibrils. Abundance of PLA dots resulting from the proximity between Aβ and (**C1**) syndapin-2 and (**C2**) Rab5 in BECs treated with Aβ assemblies at 100 nM for 15 minutes. Violin plot of *n* = 30 images. **** *P* <0.0001, one-way ANOVA comparing oligomers *versus* monomers and fibrils. (**D**) LRP1 expression in BECs treated with Aβ monomers, oligomers, and fibrils at 100 nM for 2 hours measured by Western blotting. Data was normalised to the loading control (GAPDH). LRP1 expression is normalised to untreated cells. Mean ± SD (*n* = 3). * *P* <0.05, one-way ANOVA comparing Aβ monomers and oligomers *versus* fibrils.

## Discussion

In AD, Aβ pathology progresses in a spatiotemporal pattern through the connected brain structures, involving accumulation of neurotoxic Aβ in the blood vessels and in brain parenchyma (6, 12). At the BBB, LRP1 acts as a main transporter for Aβ (11, 14, 15, 17). It is thus safe to assume that decline in the expression of LRP1, reported with normal ageing and AD (24–26), results in a deficient Aβ clearance across the BBB. More importantly, mounting evidence suggest that, apart from the depletion of LRP1 in brain blood vessels, dysfunctions in the endosomal trafficking contribute to impaired transport of Aβ (28, 32, 34, 60). Recently, we confirmed that aside from the classical vesicular endocytosis, LRP1 is also trafficked through tubular carriers across BECs, a process which is facilitated by syndapin-2, and that the avidity of the LRP1 ligand affects the overall intracellular trafficking across the BBB (40). Hence, given the importance of LRP1 and its associated Aβ-binding proteins in the clearance of Aβ and pathophysiology of AD, we investigated whether syndapin-2 is also involved in Aβ clearance from the brain through BECs, and if syndapin-2 mediated transport is altered in healthy ageing and AD. Additionally, we assessed the effect of the avidity of Aβ assemblies (monomers, oligomers, and fibrils) in their traffic across BECs.

Initially, we characterised the mechanism underlying syndapin-2-mediated LRP1 transcytosis across the BECs. We found that syndapin-2 is present as tubular structures within BECs and blood vessels in the hippocampus of WT mouse brains, where syndapin-2 interacts with LRP1 (**Fig. 1**). The tubular structures showed an estimated diameter of ~ 500 nm and lengths up to ~ 2 μm, which is in accordance to the structures observed by STED in mouse brain capillaries (40). At the molecular level, we uncovered that syndapin-2 is not associated with LRP1-related vesicular endocytic proteins and that depletion of syndapin-2 levels triggers a considerable increase in the association of LRP1 with classical vesicular endocytic proteins in BECs (**Fig. 2**). Syndapin-2 has been reported to be involved in clathrin-mediated endocytosis (61, 62), biogenesis of caveolae and caveolae endocytosis (50, 63), and clathrin- and caveolae-independent mechanisms of endocytosis (64). Our data demonstrated that syndapin-2 colocalises with clathrin within BECs, which may suggest that syndapin-2-mediated tubulation at the BBB requires clathrin for the initial deformation of the cell membrane. However, the knockdown of syndapin-2 in the BECs prompted an increase in the association of LRP1 with clathrin and endosomal proteins, indicating that, with a depletion of syndapin-2, the clathrin-mediated endocytosis is triggered to compensate for lack of the tubular mechanism. Thus, it remains to be fully elucidated the role of clathrin in the initiation of syndapin-2-mediated transcytosis of LRP1 and whether this mechanism of tubulation is dependent on clathrin or other endocytic mechanisms. Apart from the interaction with clathrin, dynamin-2 and β-actin, syndapin-2 showed a negligible association with the vesicular endocytic elements (early and late endosomes) leading us to hypothesise that the syndapin-2-mediated pathway is independent from the vesicular endocytic pathway. Our results also showed that, although tubulation-mediated transport of LRP1 is independent of the vesicular endosomal pathway, there is a balance between these two mechanisms with BECs compensating the lack of synapin-2 with an increase of endosomal proteins to drive trafficking of LRP1. Therefore, it is imperative to further understand this vesicular/tubular trafficking at the BECs and which stimulus favour either of these mechanisms in health and disease.

We then demonstrated that Aβ is closely associated with syndapin-2 in brain blood vessels (**Fig. 3, 4**) and, more importantly, we found that downregulation of syndapin-2 impairs brain-to-blood transport of Aβ and leads to higher levels of Aβ in the brain (**Fig. 3**). In neurons, it has been suggested that early and late endosomes (Rab5 and Rab7) regulate Aβ endosomal trafficking (65). However, at the BBB, early and recycling endosomes along with PICALM are critical mediators for the transport of Aβ (28). In this vesicular endosomal pathway associated with PICALM, the Aβ-LRP1 complexes are colocalised with Rab5 and EEA-1-positive early endosomes but not with Rab7, a GTPase that directs fusion of late endosomes with lysosomes, or the lysosomal-specific proteins. Rather, Aβ-LRP1/PICALM colocalised with Rab11, GTPase that regulates recycling of vesicles controlling transcytosis (66), and an inhibition or mutation of Rab11 inhibits the basal-to-apical transport of Aβ (28). Furthermore, knockdown of PICALM inhibited Rab5 and Rab11 GTPase activity in BECs treated with Aβ, indicating that PICALM binding to Rab5 and Rab11 is critical for maintaining these GTPases active during endosomal trafficking of Aβ. Based on our *in vitro* data showing that syndapin-2 is independent from vesicular endocytic elements and colocalisation of syndapin-2/Aβ in, we propose that Aβ is also transported through syndapin-2 tubular carriers across BECs. Indeed, horizontal orthogonal views obtained from our *in vitro* BBB model depicted the syndapin-2/Aβ complexes as elongated tubular-like structures not only at the basal but also apical side of BECs.

Once we established the association of LRP1/syndapin-2 and Aβ, we assessed the levels of syndapin-2 in an amyloidosis mouse model (APP-PS1). Notably, the expression levels of syndapin-2 were significantly reduced in brain blood vessels of 12-months old AD mouse brains compared to age-matched littermates, whereas early endocytic proteins, including clathrin and Rab5, were increased (**Fig. 4**). An imbalance in the levels of syndapin-2 and early endosomes was also observed in healthy ageing in WT and APP-PS1 brain blood vessels (**Fig. 5**). Importantly, age-induced loss of syndapin-2 within the brain was much greater in our genetically modified AD mice, indicating that a genetic predisposition of AD accelerates the loss of syndapin-2. Collectively, these data revealed that syndapin-2 levels are affected in healthy ageing and AD, which then may trigger LRP1/Aβ trafficking through the vesicular endosomal pathway. Interestingly, these results mirror our *in vitro* and *ex vivo* data with the syndapin-2 KO mouse brains, in which we verified that depletion of syndapin-2 triggers an increased association of LRP1 and vesicular endocytic proteins in BECs and greater Rab5 expression levels in syndapin-2 KO brains. The compensatory increased expression of Rab5 in the APP-PS1 mice brains suggests that the Aβ transport across the brain endothelium would still proceed through the vesicular pathway. Nevertheless, with an increased Aβ production and the lack of syndapin-2 to facilitate Aβ clearance, the vesicular endosomal system is, possibly, overloaded causing abnormalities in the endosomal pathway. Such overload may be reflected in the abnormalities found in early endosomes, i.e., increased number of enlarged abnormal endosomes accumulating Aβ (34–37). In addition, genetic alterations in vesicular endocytic elements might impair Aβ clearance. A reduction in PICALM levels is well-established, which in turn results in a diminished activation of Rab5 and Rab11 for endosomal trafficking (28). Hence, an enrichment of Rab5-positive endosomes may not necessarily result in a greater rate of transcytosis of Aβ as Rab5-mediated endocytosis is defective in AD. A number of studies has pointed out abnormalities in the vesicular-mediated pathway in early and sporadic AD (34–36, 38, 39). Studies of human donor tissue by Cataldo *et al.* (34, 35) have revealed that at the earliest stages of AD, many neurons have increased levels of Aβ and exhibit an abnormal overactivation of Rab5-positive early endosomes. These enlarged early endosomes exhibited immunoreactivity for other early endosomal markers (such as EEA-1 and Rab4) and Aβ, implying that accumulation of Aβ in neurons is correlated with abnormal endosomes. Interestingly, in our study using APP-PS1 mouse brains, we describe the increase in the levels of Rab5 early endosomes, however the levels of late and recycling endosomes remain unaltered suggesting that the upregulation in endosomal levels is not compensated in the later stages of endocytosis. Additionally, apart from PICALM, other studies have also identified alterations in the expression of Rab5-associated endocytic proteins, including BIN1 (31, 60) and RIN3 (32), in AD. It remains unclear how BIN1, RIN3 and other elements interact with Rab5-mediated pathway in the BECs. Nevertheless, it is conceivable to hypothesise that alterations in these elements may also contribute to abnormalities in the endosomal machinery that, ultimately, affect the trafficking of Aβ via Rab5-mediated endocytosis. Here, we demonstrated a possible role of syndapin-2 in the overactivation of Rab5-positive endosomes observed in early stages of neuropathology of AD. Hence, we demonstrate here that, apart from the endosomal dysfunctions observed in neurons, dysfunctions in the endosomal pathway are present at the blood vessels in ageing and AD, and that possibly these are a result from the downregulation of syndapin-2. It is worth to highlight that, even though the syndapin-2 levels are changed in microvessels and parenchyma of healthy and AD mouse brains, these alterations were more pronounced in the microvessels implying a prominent role of syndapin-2 in the clearance of Aβ across the brain endothelium.

Based on our previous study, we also assessed whether the avidity of different Aβ assemblies impacts the intracellular trafficking across BECs (**Fig. 6**). Even though we observed an increase in the association of LRP1/syndapin-2 in BECs treated with all different species of Aβ, we observed distinct pathways for the transcytosis of LRP1/Aβ complexes. Aβ oligomers appear to favour the formation of syndapin-2 tubular carriers for a fast shuttling across the BECs, while fibrils bias LRP1 via endosomal sorting and lysosomal sorting. Collectively, our data provides the first demonstration of the participation of syndapin-2 in LRP1-mediated Aβ clearance across BECs, and its role in the accumulation of Aβ within the brain in AD and ageing. Furthermore, we establish that the size and thus avidity of distinct Aβ assemblies plays a role in their transport across the BBB, and in expression levels of LRP1 in BECs. Hence, although the trafficking of Aβ species across BECs remains uncharted, we provide the evidence for its relevance in the clearance of Aβ. Further studies focusing on these species and their intracellular trafficking are of paramount to fully understand the faulty clearance of Aβ and develop targeted therapies for the removal of Aβ from the brain.

## Materials and Methods

### Materials

bEnd3 (CRL-2299) and Dulbecco’s Modified Eagle’s Medium (DMEM) were obtained from ATCC. Foetal bovine serum (FBS), penicillin/streptomycin, phosphate-buffered saline (PBS at pH 7.4), 0.25% trypsin-EDTA solution and rat tail collagen I were obtained from Sigma-Aldrich. Polybrene, syndapin-2 shRNA and control shRNA lentiviral particles were obtained from Santa Cruz Biotechnology. The transwell permeable support polyester membranes (0.4 μm, 1.12 cm^2^) were obtained from Corning Inc. Paraformaldehyde (PFA), Triton X-100, normal horse serum, FITC-conjugated lectin, PLA probes (anti-rabbit PLUS and anti-mouse MINUS), Duolink detection reagent orange, radioimmunoprecipitation (RIPA) buffer, Tween-20, dextran (60-76 kDa), bovine serum albumin (BSA) and thioflavin T (ThT) were also obtained from Sigma-Aldrich. Protease inhibitors, BCA protein assay kit and Laemmli sample buffer were purchased from Biorad. 5-Fluorescein-amyloid-β protein (Aβ 1-40) (4090152) and amyloid-β protein (Aβ 1-40) (4014442) were obtained from Bachem. Aβ (1-40) mouse ELISA kit, 1-404’,6-diamidino-2-phenylindole (DAPI), and Leica standard immersion oil were obtained from Thermo Fisher Scientific. Vectashield Mounting Media was obtained from Vector Labs. RealTime-Glo™ MT Cell Viability Assay was purchased from Promega. Uranyless was obtained from Delta Microscopies. Antibodies used are listed in **Table S1** in Supporting Information.

### Animals

Syndapin-2 knockout (KO) mice (6-months old) were used to investigate the role of syndapin-2 in Aβ in accumulation in the brain. Homologous recombination and the Cre-*loxP* recombination were used to generate the KO mice with a *loxP*-flanked (floxed) expression of the PACSIN2 gene, according to the previously published work (67). Wild-type (WT) C57BL/6 mice were used as a control. All procedures involving animal experiments were approved by the Animal Welfare and Ethical Review Body of the University of Portsmouth and performed by a personal license holder under a Home Office-issued project licence in accordance with the Animals (Scientific Procedures) Act 1986 (UK). Male C57BL/6J mice were used to investigate the native association of syndapin-2 with Aβ. APP-PS1 transgenic mouse model of AD (52), which carries mutations for APP and presenilin-1 (APPswe and PSEN1dE9, respectively) resulting in increased Aβ production, was used to investigate the expression levels of syndapin-2 in an amyloidosis mouse model. This line was kept by crossing transgenic APP-PS1 with C57BL/6J WT mice. In all experiments, the WT littermates were used as controls for APP-PS1 animals. 4- and 12-months old WT and APP-PS1 animals were used to assess the effect of ageing in the expression levels of syndapin-2. All animals were bred in-house in a temperature- and humidity-controlled environment under a 12-hour light/dark cycle with free access to standard chow and water.

### Cell Culture

Mouse brain endothelial cells (bEnd3) were used between passage 20-30. bEnd3 were grown in DMEM supplemented with 10% (v/v) FBS, and 100 IUL mL^−1^ penicillin/100 mg mL^−1^ streptomycin. Short hairpin RNA (shRNA) lentiviral particles were used to generate a stable cell line expressing lower levels of syndapin-2 (40). Briefly, bEnd3 cells were seeded onto a 6-well plate at a density of 100,000 cells per well and grown overnight. At 50% of confluence, cells were treated with the shRNA lentiviral particles in DMEM supplemented with polybrene (5 μg mL^−1^), and further incubated overnight. On the next day, media was replaced, and cells were maintained for 2 days. Stable clones expressing shRNA were selected by puromycin (5 μg mL^−1^). Syndapin-2 knockdown was confirmed by Western blot and immunofluorescence. bEnd3 transfected with control shRNA lentiviral particles were used as a negative control. Both syndapin-2 knockdown and shRNA control bEnd3 were cultured in DMEM supplemented with FBS, penicillin/streptomycin, and puromycin. Cells were maintained at 37 °C in an atmosphere of 5% CO2. For subculture, cells were washed with PBS, incubated with 0.25% trypsin-EDTA for 3 minutes, centrifuged and resuspended in fresh media. Media was changed every 2-3 days.

### *In Vitro* BBB Model

To obtain polarised endothelial monolayers, bEnd3 cells were seeded at 25,000 cells per cm^−2^ in collagen I-coated polyester transwells. bEnd3 cells were grown for 3 days to reach confluency, and then the media in the basal side of the Transwell was replaced to serumfree DMEM. On day 6, transendothelial resistance and permeability of dextrans (4 and 70 kDa) was measured, as previously reported (40).

### Proximity Ligation Assay

Polarised bEnd3 were washed twice with PBS, fixed in 4% (w/v) PFA in PBS for 15 minutes and permeabilised with 0.1% (w/v) Triton X-100 in PBS for 10 minutes. Proximity ligation assay (PLA) was carried out by using Duolink probes and detection reagents according to the supplier’s instructions. Briefly, monolayers were incubated with Duolink blocking solution for 1 hour at 37 °C and then incubated with the two antibodies targeting each protein of interest overnight at 4 °C. Subsequently, cells were incubated with Duolink PLA probes (anti-rabbit and anti-mouse) for 1 hour at 37 °C, washed and incubated with Duolink ligase and polymerase enzymes for 30 and 100 minutes, respectively. As a negative control, PLA protocol was followed with the exception of the addition of the primary antibodies to determine the specificity of the PLA probes. Nuclei were counterstained by incubation with DAPI for 10 minutes. Membranes were mounted in glass coverslips using Vectashield Mounting Media. Images were acquired using a Leica TCS SP8 confocal microscope, via sequential scan to reduce fluorophore bleed-through, with a 63x oil immersion objective. Z-stacks (10-20 stacks per sample) were collected (20 optical sections) and fluorescence intensity of the PLA signal was quantified using ImageJ. Abundance of PLA signal was normalised by the number of nuclei in each z-stack image. List of antibodies in Supporting Information.

### Immunofluorescence

Polarised bEnd3 were washed twice with PBS, fixed in 4% (w/v) PFA in PBS for 15 minutes, permeabilised with 0.1% (w/v) Triton X-100 in PBS for 10 minutes and incubated with 5% (w/v) BSA in PBS for 1 hour at room temperature. Subsequently, bEnd3 cell monolayers were incubated with primary antibodies diluted in 1% (w/v) BSA and 0.01% (w/v) Triton X-100 in PBS overnight at 4 °C, washed with PBS and incubated with the corresponding secondary antibody at room temperature for 2 hours. Nuclei were counterstained by incubation with DAPI for 10 minutes. Transwell membranes were excised and mounted on glass coverslips with Vectashield Mounting Media. All images were acquired at room temperature using a Leica TCS SP8 confocal microscope using a 63x oil immersion objective (z-stacks of 20 optical sections). Images were processed on ImageJ. List of antibodies in Supporting Information. Adult WT and APP-PS1 mice were anesthetised with isoflurane and pentobarbitone (1.25 mg Kg^−1^ of body weight, intraperitoneal) and transcardially perfused using fixative containing 1% (w/v) PFA and 15% (v/v) saturated picric acid in 0.1 M phosphate buffer (PB, pH 7.4), according to previously reported protocols (68). After perfusion, brains were dissected from the skull, post-fixed overnight at room temperature in fixative solution, sectioned with a vibratome and stored in a solution containing 0.1 M PB and 0.05% (v/v) sodium azide until further use. Coronal sections of the hippocampal region of WT and APP-PS1 mice brains were incubated in 20% (v/v) normal horse serum containing 0.3% (w/v) Triton X-100 in PBS for 2 hours at room temperature under gentle agitation. Then, the tissue sections were incubated with the primary antibodies diluted in 0.3% (w/v) Triton X-100 in PBS overnight at 4 °C. Following primary antibodies, sections were washed with PBS and incubated with the appropriate secondary antibodies and FITC-conjugated lectin (1:200) for 2 hours at room temperature under agitation. Tissue sections were washed with PBS and mounted on glass coverslips with Vectashield Mounting Media. Images were acquired with a Leica TCS SP8 confocal microscope using a 20x objective in a sequential mode to reduce fluorophore bleed-through. Z-stacks were acquired (20 optical sections) with images being processed on ImageJ or Leica LAS-X software for rendering 3D reconstructions.

### *In Vitro* Permeability of Aβ (1-40)

To evaluate the permeability of Aβ across the polarised bEnd3 monolayers, FAM-Aβ (1-40) (500 nM) was added to the basal or apical side of the transwell to measure basal-to-apical or apical-to-basal permeability, respectively. Samples were collected either from the apical or basal side, and fresh medium was added to replace the volume. FAM-Aβ (1-40) fluorescence was measured in a 96-well plate using a Spark Multimode microplate reader (Tecan). Apparent permeability was calculated as:

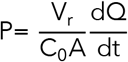

With V_r_ being the volume of the receptor (apical), C_0_ initial Aβ concentration, A the total surface area of the Transwell membrane, and 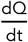 the transport rate calculated as the gradient of mass over time.

### Western Blot

Polarised bEnd3 were washed twice with PBS and RIPA buffer containing protease inhibitors (1:50) was added directly to the transwell membranes and left on ice for 1 hour. Cells were collected, centrifuged, and the supernatant was collected for Western blot analysis. WT, KO and APP-PS1 mice were anesthetised, decapitated and the brains were carefully collected from the skull. Brains were homogenised in a 4-fold excess volume of ice-cold buffer solution: 10 mM HEPES, 141 mM NaCl, 4 mM KCl, 2.8 mM CaCl_2_, 1 mM MgSO_4_, 1 mM NaH_2_PO_4_ and 10 mM glucose (pH 7.4) in a glass homogeniser (10 to 15 strokes). Microvessels and capillary-depleted fractions were prepared from WT, KO and APP-PS1 mice brains, as previously described (28). Homogenate was suspended in an equal volume of 26% (w/v) dextran (64-76 kDa), mixed, and centrifuged at 15,800 g for 10 minutes at 4 °C. Pellet containing the brain microvessels was carefully separated from the supernatant containing capillary-depleted brain (parenchyma). Both fractions, microvessels and parenchyma, were washed twice in ice-cold buffered solution by centrifugation at 15,800 g for 10 minutes. Brain microvessels and parenchyma fractions were resuspended in RIPA lysis buffer containing protease inhibitors, centrifuged for 20 minutes, and the supernatant used for Western blot analysis. Protein levels in the lysates were determined using a BCA Protein Assay Kit. Lysates were mixed with Laemmli sample buffer, and proteins (25 μg) were separated on 10% SDS polyacrylamide gels and transferred to polyvinylidene difluoride (PVDF) membranes. All membranes were blocked with 5% (w/v) non-fat milk in Tris-buffered saline (TBS) containing 0.1% (w/v) Tween-20 (TBS-T) for 1 hour and then incubated at 4 °C with the primary antibody overnight. After washing with TBS-T three times for 5 minutes, membranes were incubated with the corresponding secondary antibody for 1 hour at room temperature, washed with TBS-T and imaged using an Odyssey CLx (LI-COR Biosciences). All membranes were also probed for glyceraldehyde-3-phosphate dehydrogenase (GAPDH) as a loading control.

### Aβ (1-40) enzyme-linked immunosorbent assay (ELISA)

Both WT and KO mouse brains and plasma were collected and used for the quantification of Aβ (1-40) using ELISA. Briefly, ~100 mg of the brain tissue was homogenised in 8 volumes of cold 5M guanidine-HCl in 50 mM Tris using a glass homogeniser. Homogenates were left on an orbital shaker at room temperature for 3-4 hours. Then, samples were diluted (1:5) with cold PBS containing protease inhibitor cocktail and centrifuged at 16,000 g for 20 minutes at 4 °C. Blood was collected in heparin-coated tubes and spun at 2000 g for 10 minutes at 4 °C to remove cells from the plasma. Supernatant (plasma) was collected and diluted (1:5) with cold PBS containing protease inhibitor cocktail. Aβ (1-40) ELISA was performed according to the supplier’s instructions. Protein levels in brain and plasma homogenates were quantified using BCA Protein Assay.

### Preparation and characterisation of Aβ assemblies

Aβ assemblies were prepared according to previously described protocols (59, 69). Briefly, Aβ (1-40) peptide was dissolved in hexafluoro-2-propanol (HFIP) at 1 mM, vortexed and bath sonicated to obtain a homogenous solution, and aliquoted to microcentrifuge tubes. HIPF was removed by overnight evaporation and, once all solvent was removed, Aβ (1-40) films were stored at −80 °C until further processed. Peptide films were then resuspended in dimethyl sulfoxide (DMSO) to 5mM, bath sonicated for 10 minutes, and vortexed for 30 s. To form the Aβ oligomers, this 5 mM Aβ solution in DMSO was diluted to 100 mM with PBS, vortexed for 15 to 30 s, and incubated at 4 °C for 24 hours. Immediately before use, the oligomeric solutions were centrifuged at 14,000 g for 10 minutes at 4 °C (to remove any fibrils that might be present), and the supernatant was diluted to the final experimental concentration. Aβ fibrils were prepared by dissolving the 5 mM Aβ solution in DMSO to 100 mM in cold 10 mM HCl, vortexed for 15 s, and incubated at 37 °C for 24 hours. Monomeric Aβ was obtained by dissolving the peptide directly in DMSO at 1 mM. Protein concentration in each Aβ preparation was confirmed by using BCA Protein Assay. Aβ assemblies were characterised by measuring ThT fluorescence and transmission electron microscopy (TEM). To evaluate the presence of fibrils, ThT assay was carried out using Aβ monomers, oligomers and fibrils diluted in PBS to 10 μM. Briefly, each Aβ preparation (20 μL, 10 μM) was incubated with ThT solution (80 μL, 50 μM) at 37 °C and then the fluorescence of ThT was measured at 450/485 nm in a 96-well plate using a Spark Multimode microplate reader (Tecan). TEM imaging was performed using a JOEL JEM-2200FS microscope equipped with a field emission gun at 200 kV, and an in-column Omega filter. TEM microscope was used in energy-filtered mode with the slit inserted to increase image contrast by collecting only elastically scattered electrons. Digital Micrograph™ software (version 3.20) was used for image acquisition and processing. Images were recorded using a direct detection camera K2 IS from Gatan working in counted mode to allow for the sensitive Aβ structures to be imaged at low electron doses without beam damage. For sample preparation, 400 mesh copper grids were glow-discharged for 40 s to render them hydrophilic. Then, 5 μL of Aβ dispersions at a concentration of ~ 0.5 mg mL^−1^ were deposited onto the grid for 1 minute. The grid was blotted with filter paper and immersed in Uranyless staining solution for 40 s for negative staining. Grid was blotted again and dried under vacuum for 1 minute.

### *In vitro* effect of Aβ assemblies in BECs

Cell metabolic activity of BECs was assessed by using the RealTime-Glo™ MT Cell Viability Assay. Cells were seeded in 96-well plates at a density of 10,000 cells per well and incubated with Aβ assemblies at concentrations ranging 0.001 to 1 μM for 1 and 24 hours. RealTime-Glo™ MT Assay was carried out according to the supplier’s instructions. To assess the intracellular trafficking of the different Aβ assemblies, BECs were treated with Aβ monomers, oligomers or fibrils at 10 μM for 15 minutes, and then cells were processed for proximity ligation assay, as described above. To evaluate the effect of Aβ assemblies in the expression of LRP1, cells were treated for 2 hours with Aβ at 10 μM and then, cells were collected and analysed by Western blotting, as described above.

### Statistical Analysis

Statistical analyses and graphical evaluations were done with Prism 8.0 (Graph-Pad Inc.). All data are represented as the mean and standard deviation (SD) unless stated otherwise. Statistical comparisons were carried out using Student’s t-test or One-way ANOVA followed by a post-hoc test. A *P* < 0.05 was considered statistically significant.

## Acknowledgments

GB thanks the ERC for the starting grant (MEViC 278793) and consolidator award (CheSSTaG 769798), the EPSRC/SomaNautix Healthcare Partnership (EP/R024723/1), EPSRC Established Career Fellowship (EP/N026322/1), EPSRC/BTG Healthcare Partnership (EP/I001697/1), and Children with Cancer UK for the research project grant (16-227). MS and DML thank Alzheimer’s Research UK South Coast Network for a pump priming grant.

## Competing Interests

The authors declare no competing financial interests.

## Supplementary Materials

**Table 1.** List of antibodies.

| **Antibody** | **Dilution** | **Supplier, Catalogue Number** |
| --- | --- | --- |
| Rabbit polyclonal to syndapin-2 | 1:400 | Abcam, ab37615 |
| Mouse monoclonal to LRP1 | 1:100 | Sigma, L2420 |
| Mouse monoclonal to LRP1 | 1:1000 | Invitrogen, 37-7600 |
| Rabbit monoclonal to LRP1 | 1:1000 | Abcam, ab92544 |
| Mouse monoclonal to GAPDH | 1:1000 | Abcam, ab8245 |
| Mouse monoclonal to ApoE | 1:200 | Novus Biologicals, NB110-60531 |
| Mouse monoclonal to RAGE | 1:100 | Santa Cruz Biotechnology, sc-365154 |
| Mouse monoclonal to p-glycoprotein | 1:100 | Invitrogen, MA1-26528 |
| Mouse monoclonal to Clathrin Heavy Chain | 1:1000 | Abcam, ab2731 |
| Rabbit polyclonal to Clathrin Heavy Chain | 1:100 | Abcam, ab21679 |
| Rabbit polyclonal to dynamin-2 | 1:100 | Abcam, ab 3357 |
| Mouse monoclonal to EEA-1 | 1:100 | Sigma, E7659 |
| Rabbit polyclonal to EEA-1 | 1:100 | Abcam, ab2900 |
| Mouse monoclonal to Rab5 | 1:400 | Sigma, R7904 |
| Rabbit polyclonal to Rab5 | 1:100 | Abcam, ab13253 |
| Rabbit polyclonal to PICALM | 1:100 | Sigma, HPA019061 |
| Mouse monoclonal to Rab7 | 1:100 | Sigma, R8879 |
| Rabbit polyclonal to Rab7 | 1:100 | Abcam, ab137029 |
| Rabbit polyclonal to Rab11 | 1:100 | Abcam, ab3612 |
| Rabbit polyclonal to LAMP-1 | 1:100 | Abcam, ab24170 |
| Rabbit polyclonal to β-actin | 1:100 | Abcam, ab8227 |
| Mouse monoclonal to Aβ | 1:1000 | Biolegend, 800708 |
| Rabbit polyclonal to claudin-5 | 1:1000 | Abcam, ab15106 |
| Rabbit polyclonal to occludin | 1:1000 | Abcam, ab216327 |
| Alexa Fluor 488 goat anti-mouse IgG | 1:500 | Biolegend, 405319 |
| Alexa Fluor 647 donkey anti-rabbit IgG | 1:500 | Biolegend, 406414 |
| Dylight 800 goat anti-mouse IgG | 1:5000 | Thermo Fisher Scientific, SA535521 |
| Dylight 800 goat anti-rabbit IgG | 1:5000 | Thermo Fisher Scientific, SA535571 |

**Figure S1.**
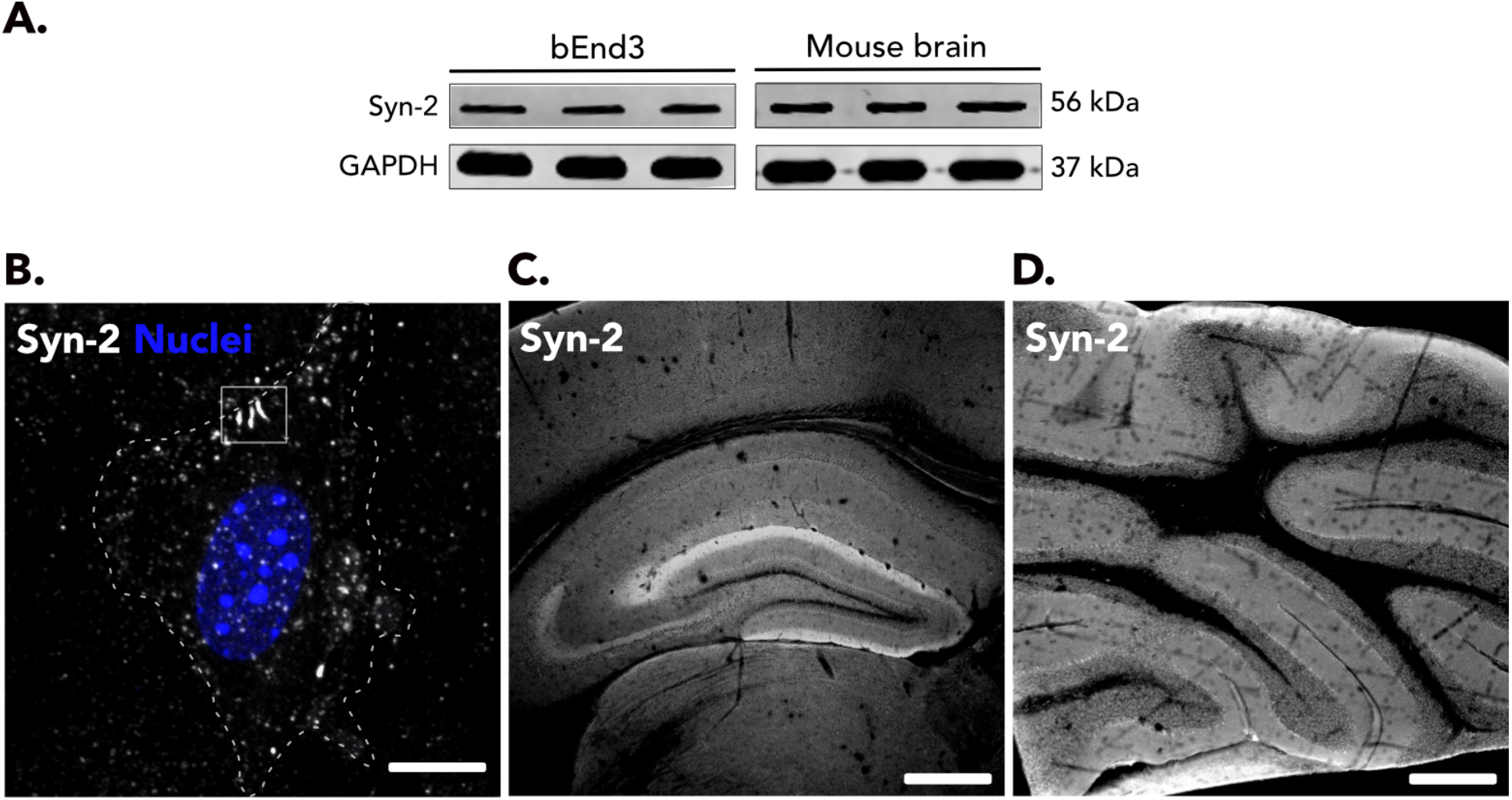
Expression of syndapin-2 in the brain endothelium and mouse brain. (**A**) Immunoblotting for syndapin-2 and GAPDH (loading control) in polarised BECs (bEnd3) and wild-type mouse brains. (**B**) Expression of syndapin-2 (in white) in BECs, highlighting syndapin-2-tubular structures. Nucleus is shown in blue and dotted line represents the cell membrane limits. Scale bar: 20 μm. Immunocolocalisation of syndapin-2 in (**C**) hippocampus and (**D**) cerebellum in mouse brain. Scale bar: 500 μm.

**Figure S2.**
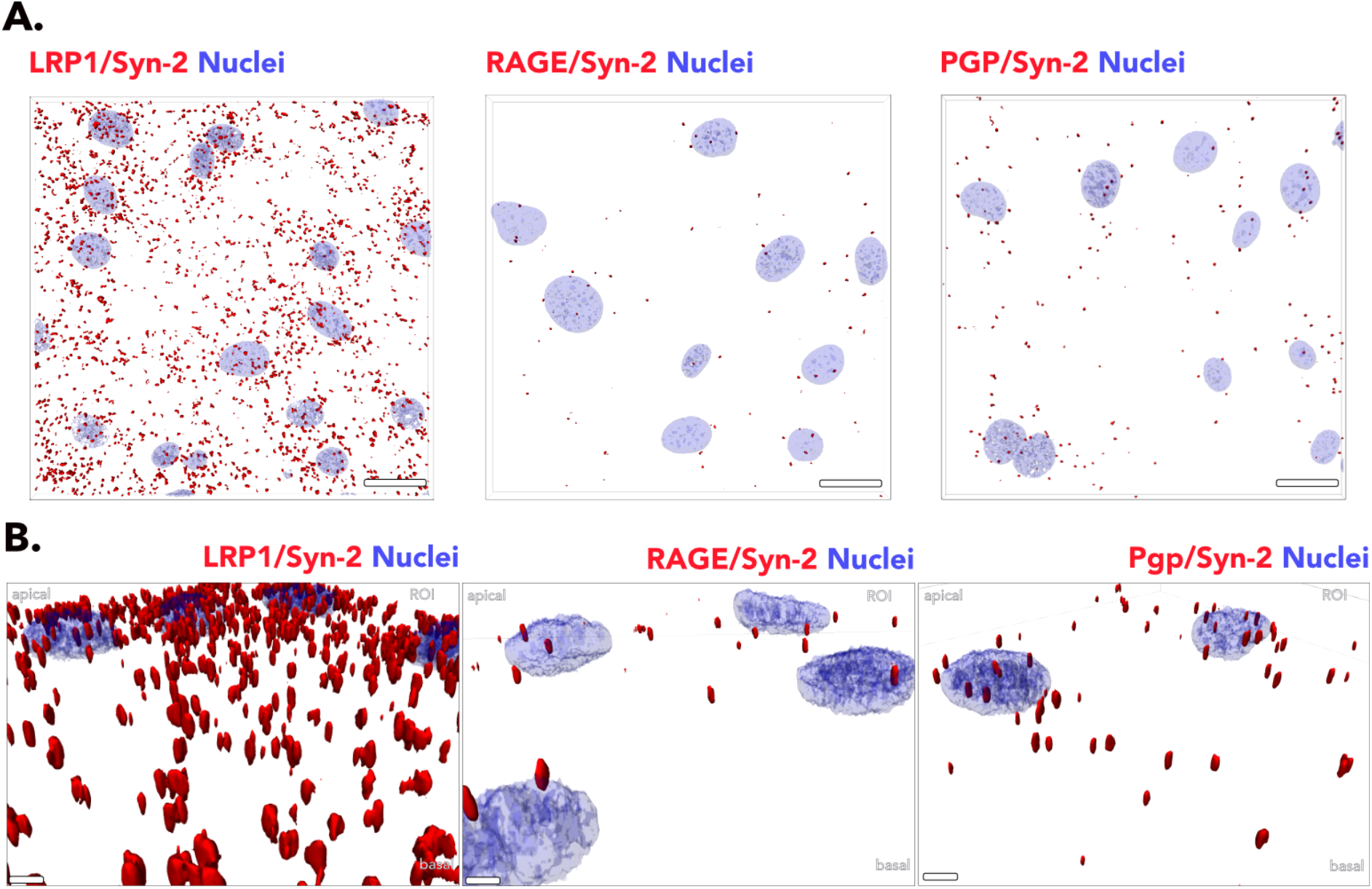
Syndapin-2 interaction with LRP1, RAGE and PGP in brain endothelium. (**A**) Representative confocal images of the proximity obtained for LRP1, RAGE or PGP with syndapin-2 in polarised BECs. Nuclei are shown in blue. Scale bar: 20 μm. (**B**) 3D renderings of BECs showing PLA dots between syndapin-2 and LRP1/RAGE/PGP. Scale bar: 2 μm.

**Figure S3.**
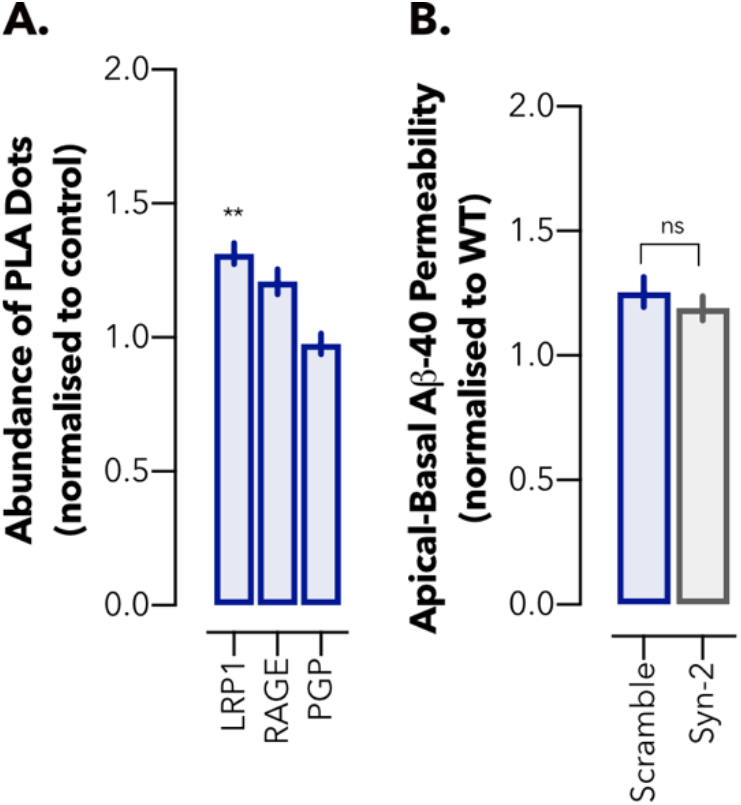
Syndapin-2 interaction with LRP1, RAGE and PGP in brain endothelium for transport of Aβ. (**A**) Abundance of PLA dots resulting from the interaction of syndapin-2 with LRP1, RAGE or PGP in the polarised BECs treated with FAM-Aβ (1-40) (500 nM) for 15 minutes. Mean ± SEM (*n* = 60 images). (**B**) *In vitro* permeability of FAM-Aβ (1-40) across shRNA control (scramble) and syndapin-2 knockdown (Syn-2) in an apical-to-basal (blood-to-brain) direction. Permeability values were normalised to wild-type (WT) BECs. Mean ± SD (*n* = 15). * *P* < 0.05, Student’s t-test.

**Figure S4.**
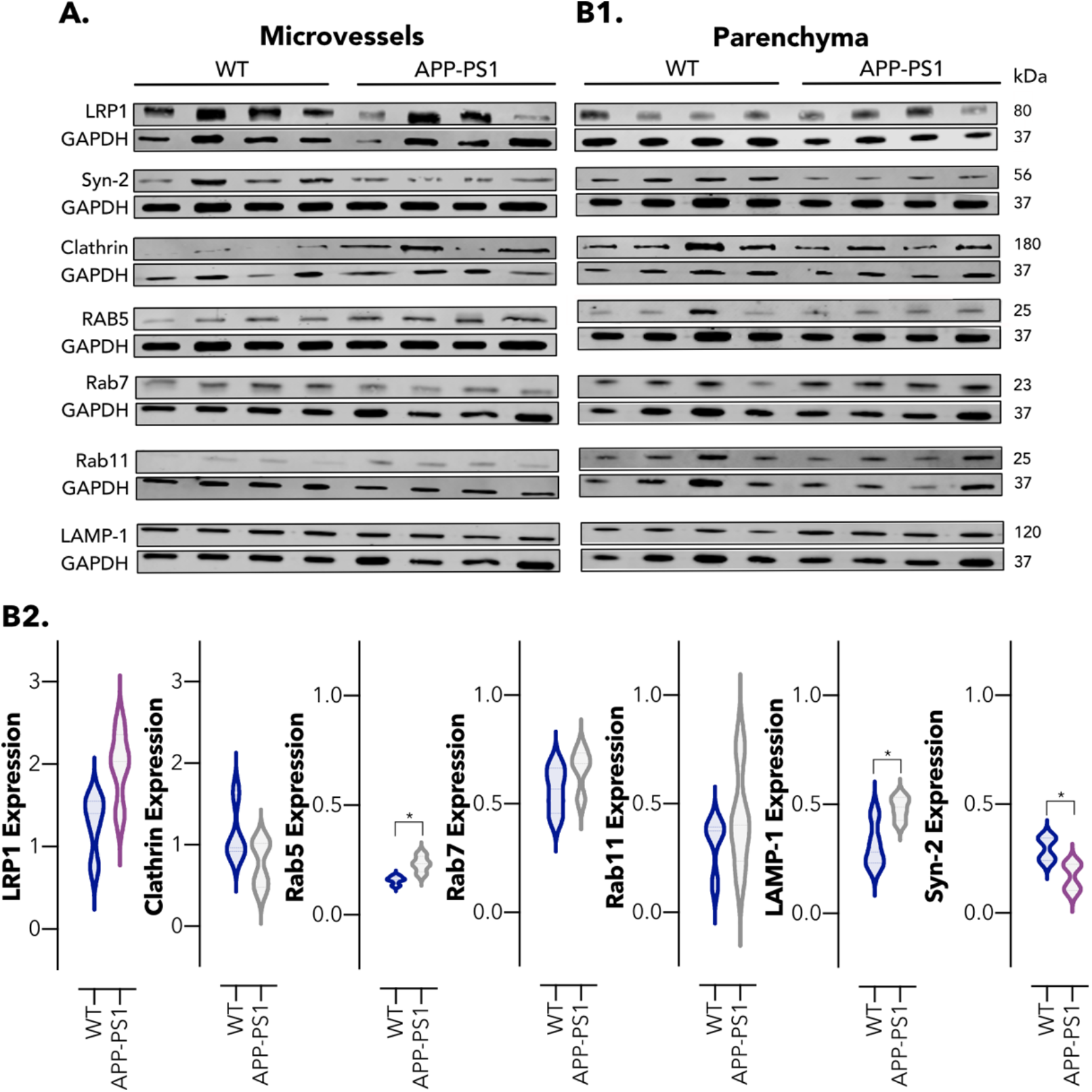
Alterations in intracellular trafficking in 12-months old WT and APP-PS1 mouse brains. Immunoblotting for endocytic proteins in (**A**) microvessel and (**B1**) parenchymal fractions of 12-months old WT and AD (APP-PS1) mouse brains. (**B2**) Relative abundance of LRP1, clathrin, Rab5, Rab7, Rab11 LAMP-1 and syndapin-2 in the parenchyma WT and APP-PS1 mouse brains. Data normalised to loading control (GAPDH). Mean ± SD (*n* = 4 animals). * *P* < 0.05, Student’s t-test.

**Figure S5.**
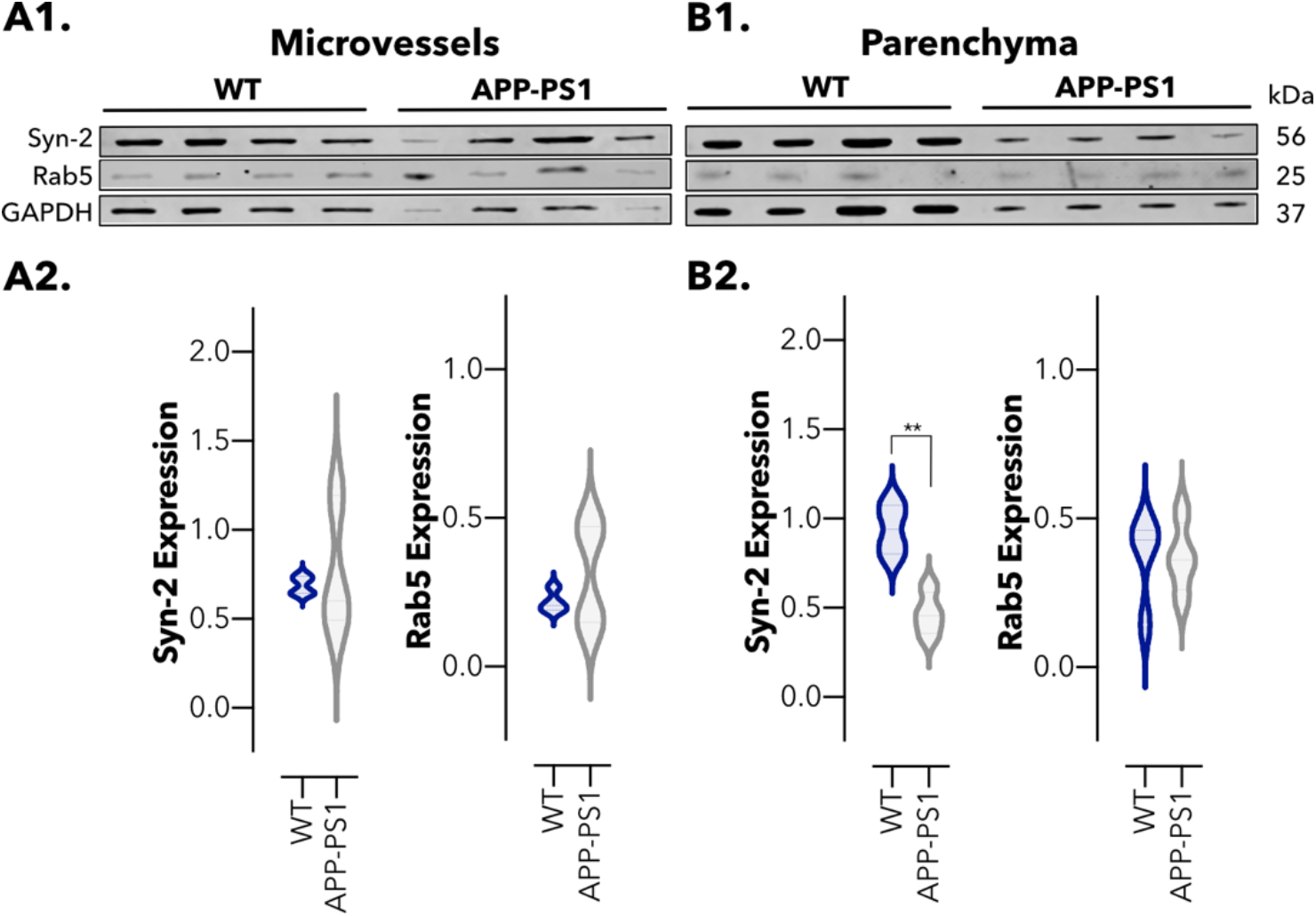
Expression of tubular and vesicular endosomal transport proteins in 4-months old WT and APP-PS1 brains. Immunoblotting for syndapin-2, Rab5 and GAPDH (loading control) in isolated (**A1**) microvessels and (**B1**) parenchyma of 4-months old WT and APP-PS1 mouse brains. Relative abundance of syndapin-2 and Rab5 in (**A2**) microvessels and (**B2**) parenchymal fractions of WT and APP-PS1 mouse brains. Data normalised to GAPDH. Mean ± SD (*n* = 4 animals). NS, non-significant, ** *P* <0.01, Student’s t-test.

**Figure S6.**
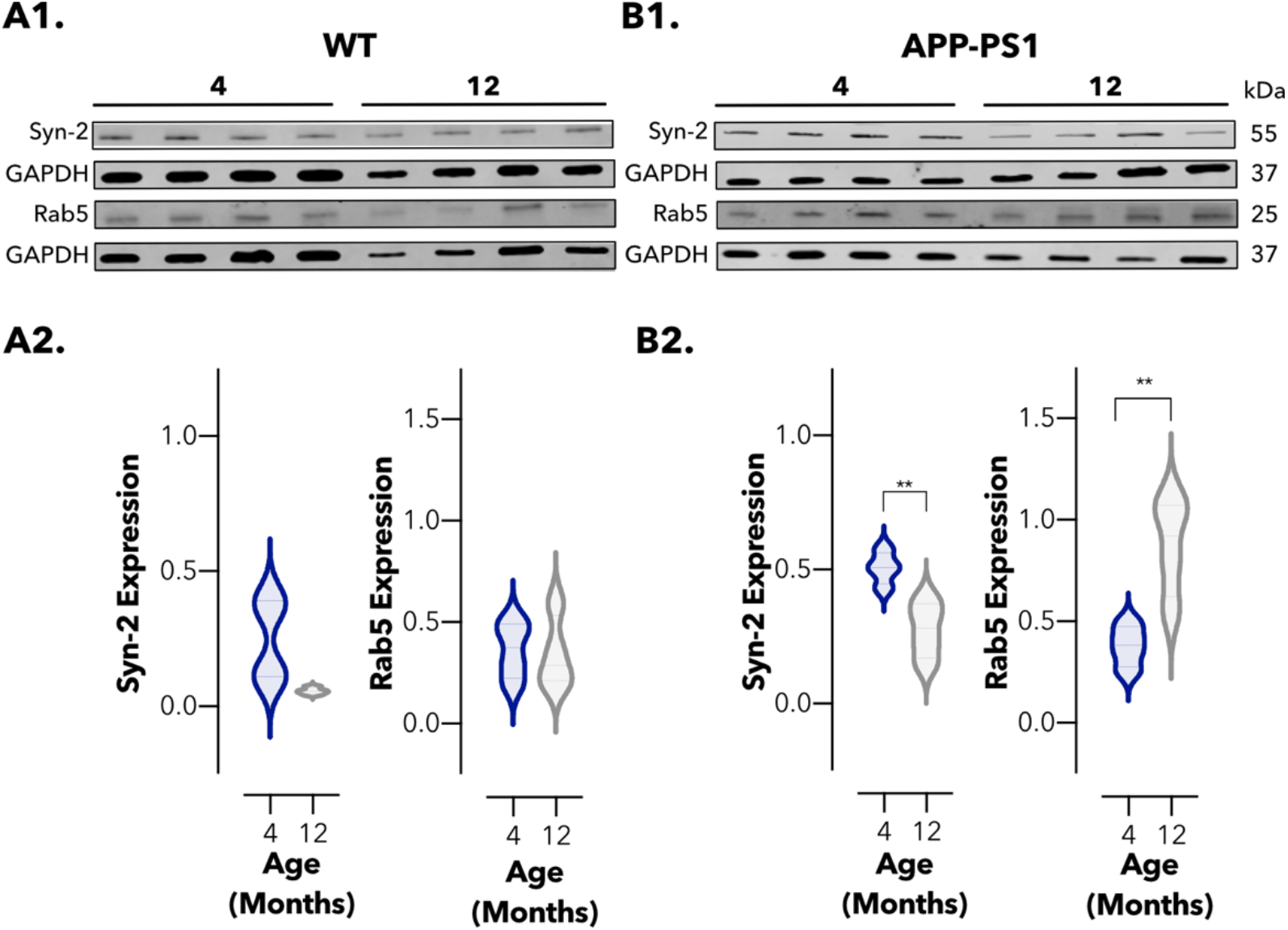
Imbalance in the level of tubular and vesicular endosomal proteins in 4- and 12-months old WT and APP-PS1 parenchyma. Immunoblotting for syndapin-2, Rab5, and GAPDH (loading control) in the parenchyma of 4- and 12-months old (**A1**) WT and (**B**1) APP-PS1 brains. Relative abundance of syndapin-2 and Rab5 levels in the parenchyma of (**A2**) WT and (**B2**) APP-PS1 brains. Data normalised to loading control (GAPDH). Mean ± SD (*n* = 4 animals). ** *P* < 0.01, Student’s t-test.

**Figure S7.**
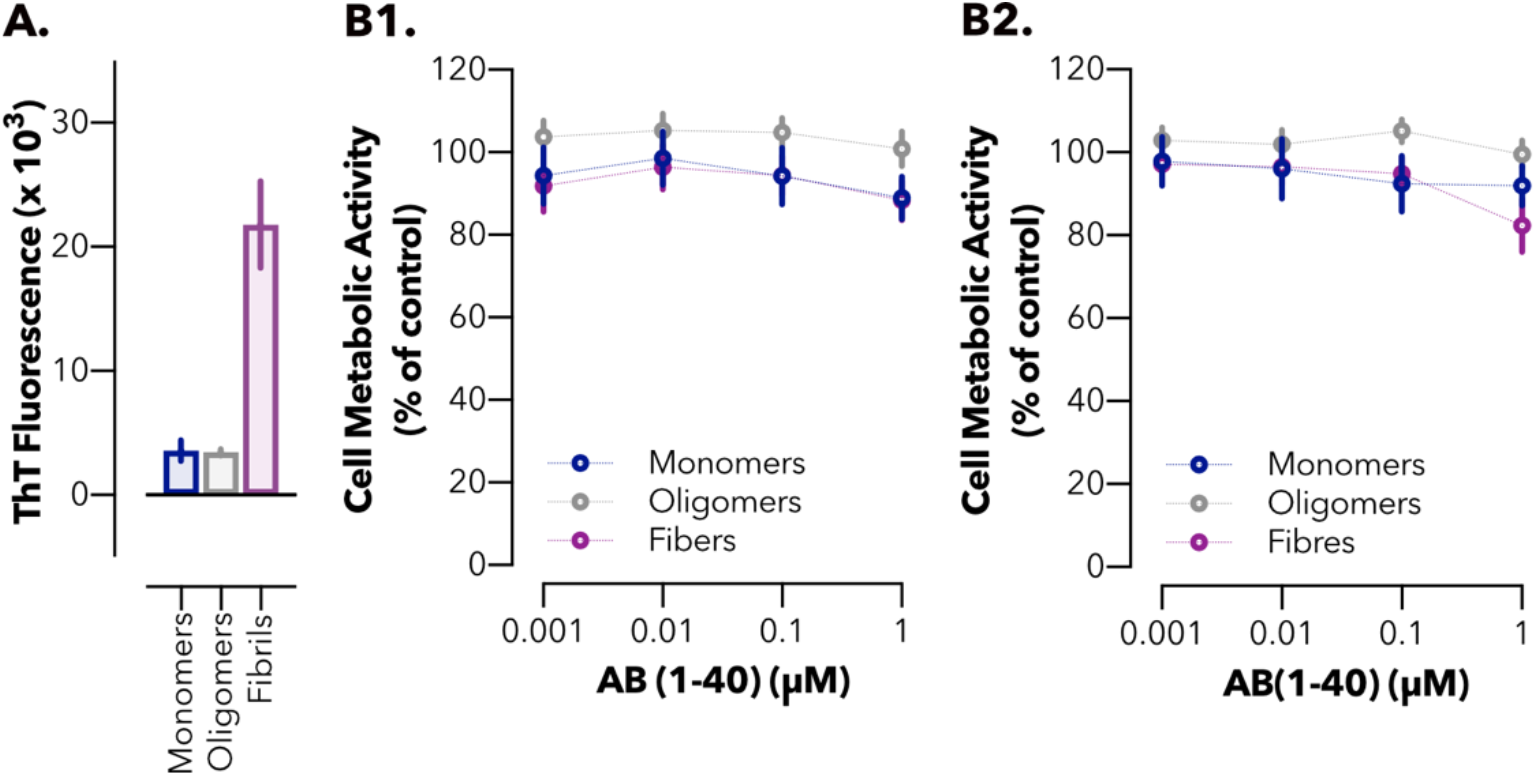
Aβ characterisation and effect on brain endothelium viability. (**A**) Thioflavin T (ThT) fluorescence intensity after incubation with Aβ monomers, oligomers and fibrils at 10 μM. Mean ± SD (*n* = 3). Cell metabolic activity of BECs treated with Aβ assemblies at concentrations ranging 0.001-1 μM for (**B1**) 1 and (**B2**) 24 hours. Mean ± SD (*n* = 12).

## Notes

### Competing Interest Statement

The authors have declared no competing interest.

### Summary of Updates

We repeated our analysis in syndapin-2 KO animals confirming our original hypothesis demonstrating that indeed down-regulation of syndapin-2 leads to amyloid-β accumulation. We also included a more detailed analysis of the transport of amyloid-β as monomer, oligomers, and fibrils, showing that their localisation and cellular sorting differs with respect to Syndapin-2.

